# Structure of the processive human Pol δ holoenzyme

**DOI:** 10.1101/872879

**Authors:** Claudia Lancey, Muhammad Tehseen, Vlad-Stefan Raducanu, Fahad Rashid, Nekane Merino, Timothy J. Ragan, Christos Savva, Manal S. Zaher, Afnan Shirbini, Francisco J. Blanco, Samir M. Hamdan, Alfredo De Biasio

**Author notes:** These authors contributed equally to the work (Claudia Lancey, Muhammad Tehseen). Correspondence should be addressed to S.M.H. or A.D.B.

## Abstract

In eukaryotes, DNA polymerase δ (Pol δ) bound to the proliferating cell nuclear antigen (PCNA) replicates the lagging strand and cooperates with flap endonuclease 1 (FEN1) to process the Okazaki fragments for their ligation. We present the high-resolution cryo-EM structure of the human processive Pol δ-DNA-PCNA complex in the absence and presence of FEN1. Pol δ is anchored to one of the three PCNA monomers through the C-terminal domain of the catalytic subunit. The catalytic core sits on top of PCNA in an open configuration while the regulatory subunits project laterally. This arrangement allows PCNA to thread and stabilize the DNA exiting the catalytic cleft and recruit FEN1 to one unoccupied monomer in a toolbelt fashion. Alternative holoenzyme conformations reveal important functional interactions that maintain PCNA orientation during synthesis. This work sheds light on the structural basis of Pol δ’s activity in replicating the human genome.

---

Three DNA polymerases (Pols), α, δ and ε replicate the genomic DNA in eukaryotes, with the latter two possessing the proofreading exonuclease activity required for high fidelity DNA synthesis^1,2^. Pol δ replicates the lagging strand and may share a role with Pol ε in replicating the leading strand ^3–10^. In contrast to the continuous leading strand synthesis, the lagging strand is synthesized discontinuously in ~200 nucleotide (nt)-long Okazaki fragments, which are then ligated to form the contiguous lagging strand^2^. Synthesis of each Okazaki fragment starts with the low fidelity Pol α synthesizing a ~30 nt RNA/DNA initiator primer. Replication factor C (RFC) then loads the homotrimeric clamp PCNA at the primer/template (P/T) junction^11^. PCNA encircles the duplex DNA and tethers Pol δ to the DNA enhancing its processivity from few nucleotides to hundreds of nucleotides per DNA binding event^12–14^. PCNA has also been shown to increase the nucleotide incorporation rate of Pol δ^15^. Additionally, the interaction of PCNA with Pol δ is critical for coordinating its transient replacement by other PCNA partner proteins. In the maturation of Okazaki fragments, Pol δ invades the previously synthesized Okazaki fragments to gradually displace the RNA-DNA primers for their removal by the PCNA-bound FEN1^16^. In translesion DNA synthesis, Pol δ is transiently replaced by a PCNA-bound translesion DNA polymerase to ensure the continuation of DNA replication^17^.

Mammalian Pol δ consists of a catalytic subunit and three regulatory subunits (Figure 1a). The catalytic subunit (p125) harbours the polymerase and exonuclease activities, and a metal-binding C-terminal domain (CTD). The regulatory subunits (p50, also referred to as the B-subunit, p66 and p12) are required for optimal activity of the holoenzyme^18^ and there is evidence of different context-specific subassemblies of Pol δ *in vivo*^19–21^. In particular, DNA damage or replication stress triggers the degradation of the p12 subunit, resulting in the formation of a three-subunit enzyme with an increased capacity for proofreading^20,21^. While mammalian Pol δ was identified more than forty years ago^22^, the architecture of this essential enzyme and its interaction with PCNA are still poorly understood. The recently published cryo-EM structure of *Saccharomyces cerevisiae* (*Sc*) heterotrimeric Pol δ bound to (P/T) DNA elucidated the interactions between the catalytic subunit (Pol3, homologous to human p125) and the two regulatory subunits (Pol31 and Pol32, homologous to human p50 and p66, respectively), showing a unique molecular arrangement^23^; the p12 subunit is absent in *Sc* Pol δ. However, how eukaryotic Pol δ achieves processive DNA synthesis and how it cooperates with PCNA and other factors during Okazaki fragment processing remains unknown.

**Figure 1.**
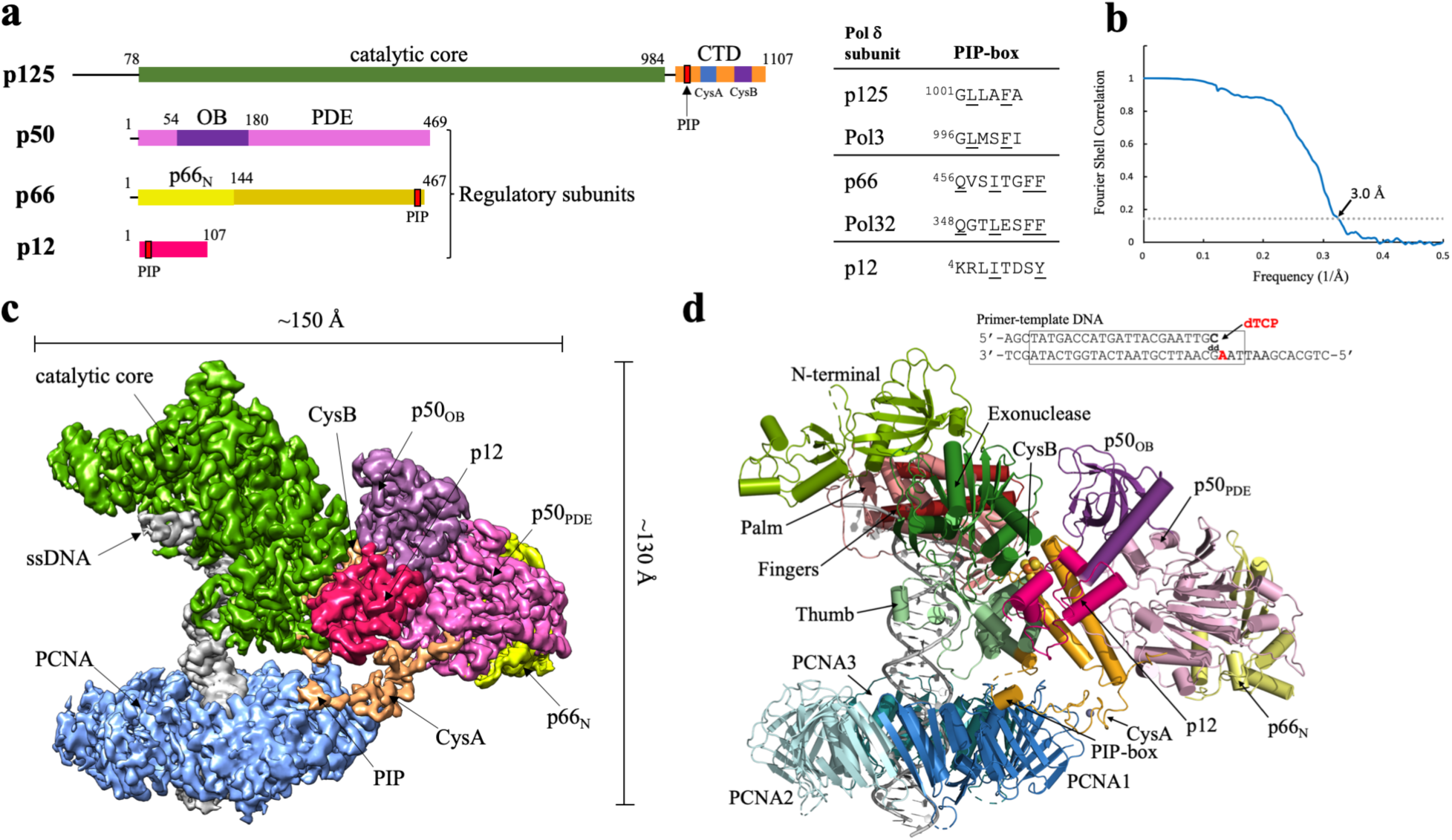
Cryo-EM structure of the processive Pol δ-DNA-PCNA complex. **a)** Domain organization of the four subunits of human Pol δ and amino acid sequence of PCNA-interacting (PIP-box) motifs. CTD: C-terminal domain; OB: oligonucleotide binding domain; PDE: phosphodiesterase domain. **b)** Gold-standard Fourier shell correlation for the Cryo-EM reconstruction of the Pol δ-DNA-PCNA complex, showing the resolution estimation using the 0.143 criterion. **c)** Cryo-EM density map of the Pol δ-DNA-PCNA complex colored by domain. **d)** Structure of the Pol δ-DNA-PCNA complex colored by domain and sequence of the DNA primer/template substrate. The region of the substrate that was modelled is boxed.

To gain insight into these molecular mechanisms, we used cryo-EM single-particle reconstruction and determined the structure of the human Pol δ heterotetramer bound to (P/T) DNA and PCNA at 3.0 Å resolution (Figure 1b). We show that this complex exists in alternative conformations where key interactions regulating the holoenzyme activity are lost. In addition, we present the structure of the Pol δ—PCNA—DNA—FEN1 complex at 4.0 Å resolution, providing the first structural evidence for the toolbelt model of PCNA function in eukaryotes. We discuss these structures in the context of Pol δ function in Okazaki fragment synthesis and maturation.

## RESULTS AND DISCUSSION

### Architecture of the processive Pol δ holoenzyme and interaction with PCNA

We firstly reconstituted a replication-competent holoenzyme comprising Pol δ, PCNA, a 25/38 (P/T) DNA bearing a 3’-dideoxy chain terminator in the primer strand and dTTP as an incoming nucleotide. A multi-subunit system was used to optimize the coexpression of human Pol δ’s heterotetrameric complex in baculovirus-infected Sf9 insect cells. We determined the cryo-EM structure of the Pol δ–PCNA–DNA–dTTP complex at 4.1 Å resolution (Supplementary Figures 1-2). Reconstitution of this complex in the presence of FEN1 followed by gel filtration (Supplementary Figure 3) led to an analogous map albeit at higher resolution (3.0 Å; Supplementary Figures 4-5) in which FEN1 is invisible, as well as to a 4.0 Å resolution map in which FEN1 is visible and could be modelled (see below in the text). We therefore discuss the structure of the Pol δ—PCNA—DNA—dTTP complex based on the 3.0 Å reconstruction (Figure 1c-d) (Supplementary Data Table 1). The structure has approximate dimensions of 150 Å × 130 Å × 100 Å and displays the catalytic domain of p125 on top of the front face of PCNA in an open configuration, while the CTD domain of p125 and the p50, p66 and p12 regulatory subunits are positioned laterally (Figure 1c-d). This arrangement allows PCNA to thread and stabilize the duplex DNA exiting the catalytic cleft. The architecture of the catalytic and p50—p66 regulatory subcomplex of Pol δ is conserved between human and yeast^23^ (RMSD_Cα_ ~ 2 Å) (Supplementary Figure 6). Importantly, our map allowed us to identify and model the p12 subunit of Pol δ (absent in yeast), showing that it bridges the exonuclease domain, the CTD domain and the oligonucleotide binding (OB) domain of the p50 subunit (Figure 1c-d). In addition, the critical region of the CTD interacting with PCNA, encompassing the C-terminus of the thumb domain and the CysA motif, was invisible in the cryo-EM map of the isolated yeast holoenzyme^23^, while it becomes visible in the human processive complex (Figure 1c and 2a). Pol δ is tethered to only one of the three PCNA monomers. Two main interaction sites are observed: a) a short parallel β-sheet involving Cys991, Thr993, Leu995 of the CTD of p125 forming four main-chain hydrogen bonds with residues Asp120 and Asp122 on the IDCL of PCNA (Figure 2, Inset 1) and b) a one-turn α-helix formed by Pol δ residues 1001-1005 which inserts into the canonical PIP-box hydrophobic cleft of PCNA through a two-fork plug made of conserved Leu1002 and Phe1005 (Figure 2, Inset 2). The CTD residues connecting the two interacting regions (residues 997-1000) are invisible, suggesting that they remain flexible. This is a mode of binding to PCNA not observed before. In particular, the newly identified p125 PIP-box (^1001^GLLAFA, Figure 1a), which highly diverges from the conserved PIP-box (with strict consensus sequence *Qxxhxxaa*, where *h* is a hydrophobic, *a* is an aromatic, and *x* is any residue), adds to the notion that PCNA can bind a diverse range of sequences broadly grouped as PIP-like motifs^24^.

**Figure 2.**
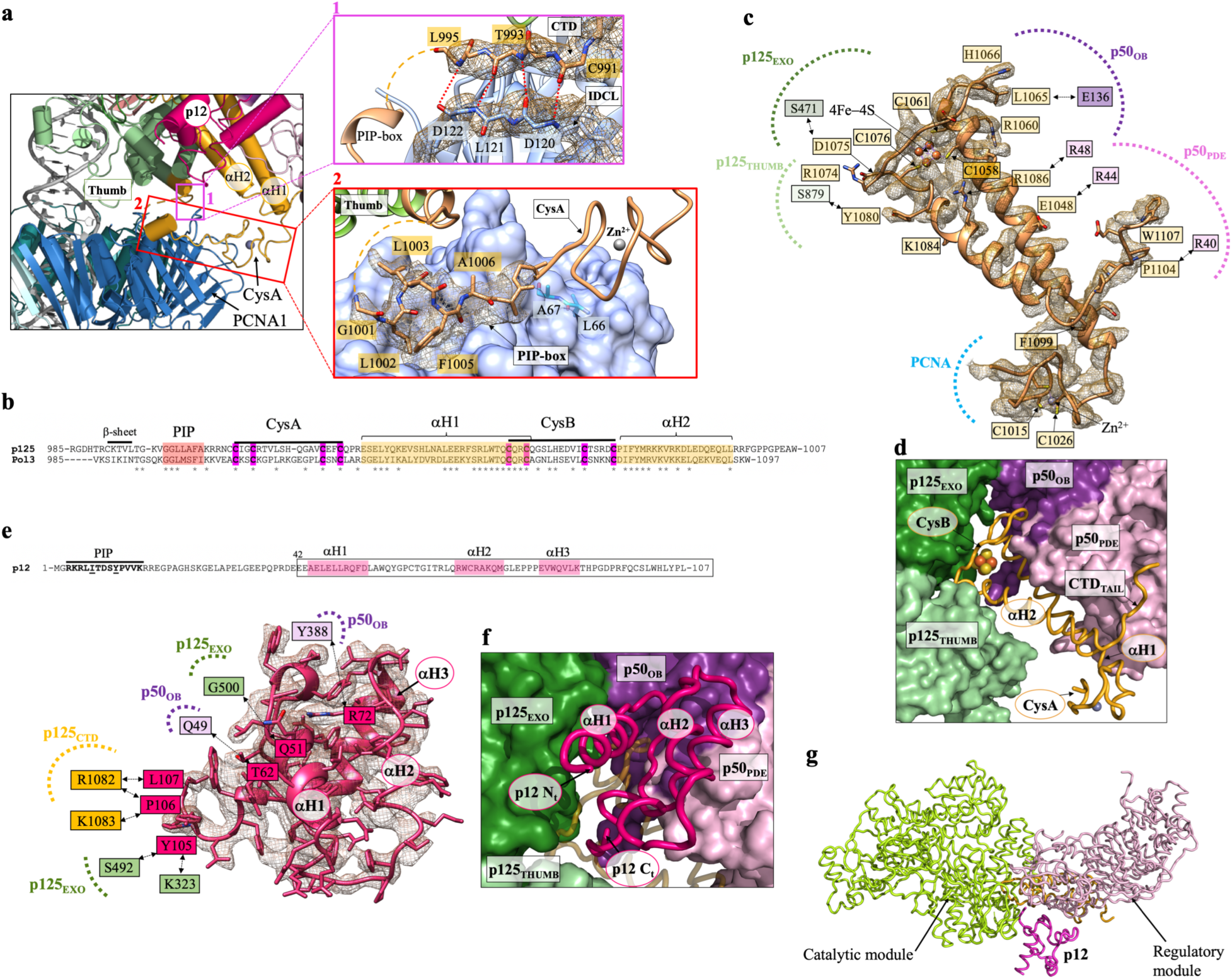
Cryo-EM density and model of selected regions of the Pol δ-DNA-PCNA complex. **a)** Map regions showing the critical interactions tethering the polymerase to PCNA. Models are colored by domain. *Inset 1*: main-chain hydrogen bonds between the CTD of p125 and the IDCL of PCNA, indicated as red dotted lines. Residues involved in the interactions are boxed and colored by domain. Amino acid side chains are not shown. *Inset 2*: interaction between the p125 PIP-box and PCNA. **b)** Sequence alignment of the CTD of human and *Saccharomyches Cerevisiae* Pol δ. Asterisks correspond to conserved residues. Conserved cysteines in CysA and CysB motifs are highlighted in magenta. **c)** Map region and model of the CTD of p125. Interfacial residues are boxed and colored by domain. Residues participating in polar interactions are connected by double-headed arrows. Some of the amino acid side chains are omitted for clarity. **d)** Model region showing the pocket between the p125 and p50 subunits where the CTD is inserted. p125 and p50 domains are shown as surfaces, and the CTD as a ribbon. The FeS cofactor and zinc ion in the CTD are shown as spheres. The p12 subunit was removed for clarity. **e)** Map region and model of the p12 subunit of Pol δ, and p12 amino acid sequence and motifs. Interfacial residues are boxed and colored by domain. Residues participating in polar interactions are connected by double-head arrows. **f)** Model region showing the position of p12 relative to the holoenzyme. p12 and CTD are shown as ribbons and the latter is shown with enhanced transparency. p125 and p50 domains are shown as surfaces. **g)** Model of human Pol δ in ribbon representation, showing the p12 subunit connecting the catalytic and regulatory modules.

**Figure 3.**
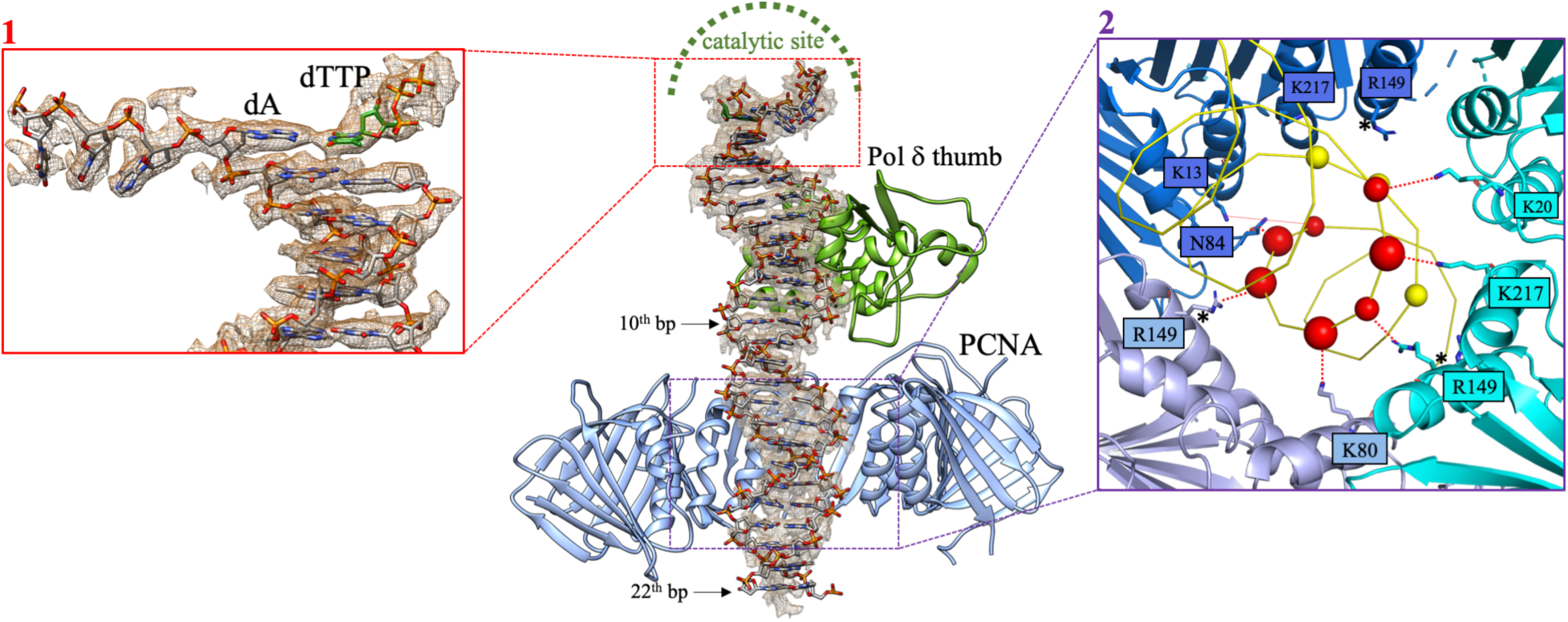
Interaction of the processive Pol δ holoenzyme with DNA. Map region around dsDNA and model, showing that DNA is held in place by the polymerase thumb domain, and stabilized by the PCNA central channel. The N-terminal, palm and fingers domains of Pol δ are removed for clarity. *Inset 1*: Cryo-EM map region and model of primer/template DNA in the Pol δ active site. The terminal adenine in the primer strand and paired incoming dTTP are labeled. *Inset 2*: PCNA interactions with DNA. PCNA subunits are shown as cartoons and colored in different shades of blue. DNA is shown as a yellow ribbon. DNA phosphates within a coulombic interaction distance (<6 Å) from PCNA residues are shown as spheres. Interacting phosphates on the template and primer strands are shown in red and yellow, respectively. Phosphates closer than 5 Å are shown with a larger sphere diameter. PCNA interacting residues are shown as sticks and labeled. The asterisk indicates that the side chain of R149 is flexible, but could approach a phosphate group at a distance <6 Å.

**Figure 4.**
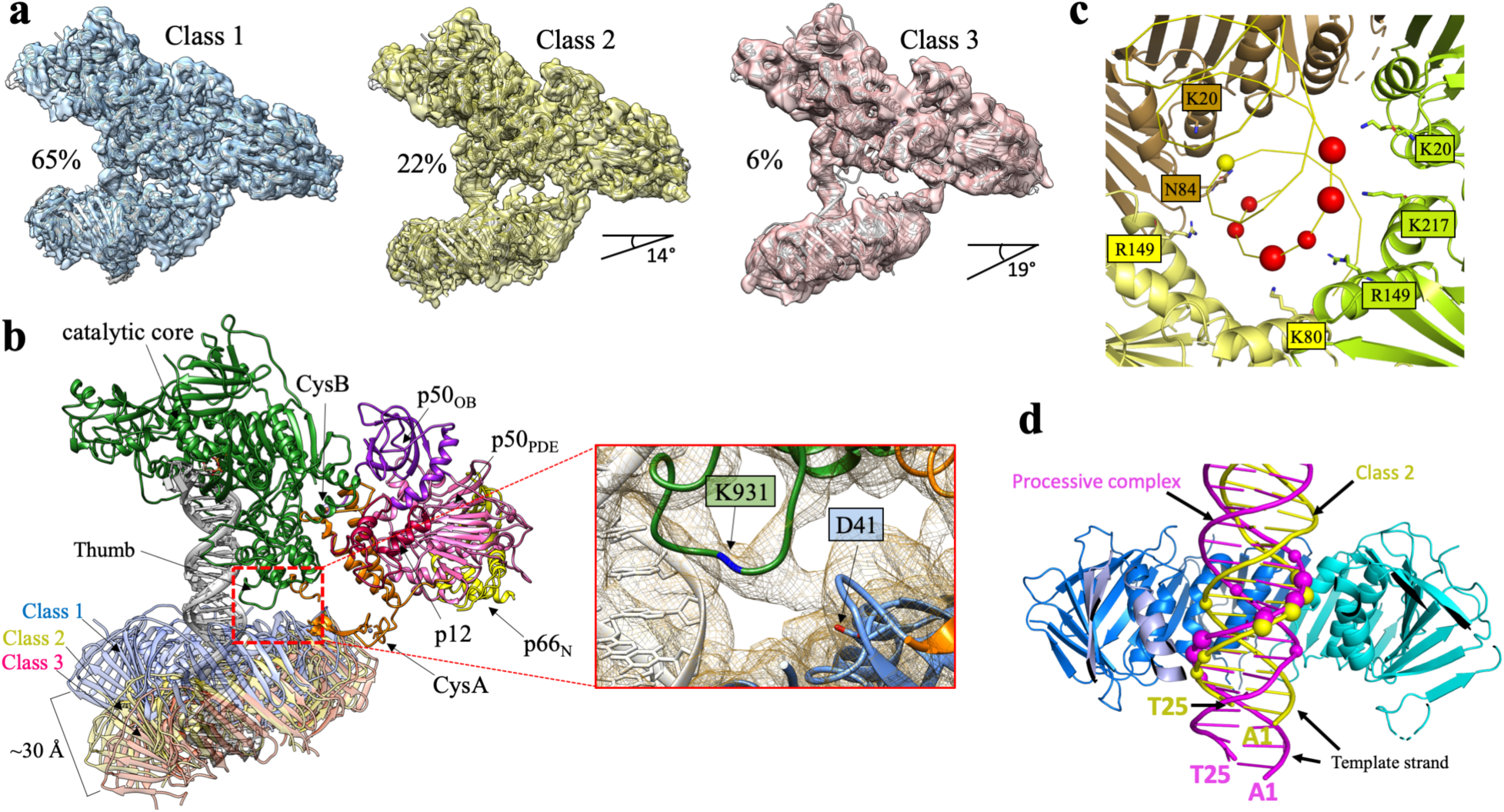
Alternative conformers of Pol δ holoenzyme. **a)** Three representative EM 3D classes showing increasing tilting of the PCNA ring relative to the polymerase. The percentage values indicated under the EM reconstructions correlate with the proportion of particles assigned to the three 3D classes and represent an estimate of the prevalence of each conformation in the data set. **b)** Overlay of the models corresponding to the EM reconstructions shown in a). *Inset*: close up of the EM map of Class 1 showing the contact between the Pol δ thumb domain and PCNA loops. Residues K931 in the Pol δ loop and D41 in the PCNA loop are labelled. Density of K931 side chain is missing, indicating that it is involved in a flexible interaction with PCNA. **c)** PCNA interactions with DNA in Class 2. PCNA subunits are shown as cartoons and colored in different shades. DNA is shown as a yellow ribbon. DNA phosphates within a coulombic interaction distance (<6 Å) from PCNA residues are shown as spheres. Interacting phosphates on the template and primer strands are shown in yellow and red, respectively. Phosphates closer than 5 Å are shown with a larger sphere diameter. PCNA interacting residues are shown as sticks and labeled. **d)** Side-view of the processive complex and Class 2 structures aligned using PCNA. The polymerase component is hidden for clarity. PCNA subunits are shown in different shades of blue and the subunit in the foreground is hidden. DNA molecules in the processive and Class 2 complexes are shown as purple and yellow ribbons, respectively. Phosphates interacting with PCNA residues are shown as spheres. Terminal bases of the DNA substrates are labelled.

We first performed a replication assay to probe the role of the interaction site involving the IDCL of PCNA on the activity of Pol δ. We showed that a PCNA variant in which the key interacting residues were simultaneously mutated (D120A, L121A, D122A and V123E) maintained the trimeric structure (Supplementary Figure 7) but severely compromised the activity of Pol δ (Supplementary Figure 8a-b), demonstrating their indispensable role for binding to Pol δ. The binding of PCNA to only one Pol δ PIP-box in our structure is a striking observation since functional assays in *Sc* Pol δ showed that three PIP-boxes located in the Pol3, Pol31 and Pol32 subunits are required to achieve processive DNA synthesis^25^. However, the putative PIP-box in Pol31 (^321^DKSLESYFNG) that is non-conserved in human, is located in a structured region, and would be positioned >40 Å away from the closest binding site on PCNA, suggesting that it may not take part in the interaction with PCNA. Besides the PIP-box on the p125 subunit, human Pol δ has two additional PIP-boxes in the p66 and p12 subunits (Figure 1a), both of which are able to interact with PCNA^26–28^. The p66 and p12 PIP-boxes are located at the extreme C- and N-terminus of the subunit, respectively (Figure 1a), and are both in flexible regions invisible in the cryo-EM map. The last visible residue of p66 is Ala144 and that of p12 is Glu42, at approximate distances of 50 Å and 36 Å from the nearest PIP-box site, respectively (Supplementary Figure 9). Therefore, both p66 and p12 PIP-boxes may possess a capture radius able to reach one or more of the PCNA binding sites. We performed activity assays to assess the role of each of the three PIP-boxes of human Pol δ (Supplementary Figure 10). We found that mutation of the key residues in the PIP-box of p125 (L1002A and F1005A) severely reduced the activity of Pol δ, while mutating the key residues in the PIP-box of p66 (F462A and F463A) or p12 (I7A and Y11A) have minimal effect (Supplementary Figure 10a). To investigate the role of the three PIP-box motifs in processivity of Pol δ, we added heparin sulphate at the start of the reaction to trap dissociated DNA polymerase molecules. We observed sever defect in the case of the PIP-box mutant of p125 followed by the PIP-box mutant of p12 and no detectable defect in the case of the PIP-box mutant of p66 (Supplementary Figure 10b). These results support our structural proposition that the PIP-box on p125 is the primary binding site to PCNA. However, the polymerase occasionally falls off the DNA during synthesis and primarily employs the long-reach PIP-box in the p12 subunit to prevent its dissociation into solution, in agreement with previous findings^21^.

The conserved CysA motif in the CTD of p125, previously reported to be important for the processivity of *Sc* Pol δ holoenzyme^29^, connects to the PIP-box and folds into a zinc finger situated next to one of the outer β-sheets of the PCNA ring (Figure 2). However, mutation of the PCNA residues closest to CysA (L66A and A67E; Figure 2, Inset 2) did not affect the trimeric structure of PCNA (Supplementary Figure 7) and the activity of Pol δ (Supplementary Figure 8a). This suggests that the CysA motif may act as an outrigger that stabilizes the holoenzyme structure and orients its interaction with PCNA rather than directly anchoring the polymerase to the clamp.

### Role of the CysB motif and p12 subunit in human Pol δ

The CysB motif in the CTD of the p125 subunit (Figure 2b), which contains an iron-sulfur (FeS) cluster^30^, is well resolved in the map (Figure 2c). We confirmed the presence of the FeS cluster in our Pol δ preparation as described previously^29,31^. A freshly purified Pol δ exhibited a yellow-brownish color and a broad absorption band that peaks at ~410 nm (Supplementary Figure 11). The extinction coefficient at 410 nm yielded a value of 13576 ± 1981 M^−1^cm^−1^ demonstrating the contribution of 3.4 ± 0.5 [Fe]/[Pol δ] (Supplementary Figure 11). The FeS cluster therefore is properly incorporated during expression and is maintained to a good extent during purification. The CysB motif contains the four conserved cysteines coordinating the FeS center and is connected to the zinc finger in the CysA motif by two antiparallel α-helices (helix α1 and helix α2), which are inserted between the catalytic domain and the p50 subunit (Figure 2c-d). Interestingly, the CTDs of Pol α and ε form equivalent complexes with their respective B-subunits but are larger and only coordinate divalent Zn^2+^ ions^32–36^. The different size of the CTD and orientation relative to the B-subunit may correlate with the distinct functional dependencies of the three polymerases on PCNA (Supplementary Figure 12); PCNA inhibits the primer extension activity of Pol α^37,38^ and modestly increases the activity of Pol ε through relatively weak interactions^39,40^. Our results confirm the important structural and functional role of the FeS cluster previously shown in both yeast and human Pol δ^29,30^. Jozwiakowski and co-workers showed that, differently from the scenario in yeast^29^, disruption of the FeS cluster does not inhibit the assembly of the human holoenzyme but rather partially destabilizes the tetrameric structure, and this effect is fully reversed by the presence of PCNA^30^. In agreement with this, our structure shows that PCNA critically stabilizes the CTD, and also suggests that the p12 subunit may stabilize the FeS-deficient holoenzyme by providing additional inter-subunit contacts (see below in the text). Loss of FeS leads to strong defects in DNA synthesis and exonucleotic activity in human Pol δ, coupled to an impaired ability to bind dsDNA^30^. This is consistent with our structure showing the FeS cluster inserting into a deep cavity between the thumb (which anchors the duplex portion of the DNA substrate) and the exonuclease domain (Figure 2d), and contacting both domains through a set of long-side chain residues forming polar and Van der Waals interactions (*e.g.*, R1074; D1075; Y1080; K1084) (Figure 2c). The [4Fe4S]^2+^ cluster in yeast Pol δ can be reversibly oxidized to [4Fe4S]^3+^ resulting in a significant slowing of DNA synthesis^41^, and this may be used by the polymerase to sense oxidative stress and stall replication under mutagenic conditions. How this DNA-mediated redox signal is transferred to the FeS cofactor (situated >25 Å away from the DNA) and how it may decrease the synthetic activity of Pol δ remains unclear. Possibly, [4Fe4S]^2+^ oxidation results in a long-range conformational change in the thumb domain which hinders the translocation of the DNA substrate through PCNA.

The CTD connects the catalytic domain of p125 to the p50-p66_N_ subcomplex, which forms an elongated prawn-like structure that projects sideways relative to p125 with no direct interaction with either PCNA or DNA (Figure 1c). The CysB motif, helix α1 and helix α2, and a loop at the C-terminus of helix α2 interact with the OB domain and the inactive phosphodiesterase domain (PDE) of p50, burying ~2200 Å^2^ at the interface (Figure 2c-d). The main interface is conserved in the human and yeast complex^23^, and the residues involved (*e.g.*, Glu1048, Arg1060 and His1066 in CysB and Glu136, Asp137 and Glu138 in p50) agree with previous biochemical and genetics studies^42,43^. However, the interaction involving the hydrophobic C-terminal tail of helix α2 (from Phe1099 to Trp1107), which inserts into a groove below p50_PDE_ (Figure 2d), is unique to the human holoenzyme, and may further stabilize the catalytic module relative to the regulatory one. Two loops in p50_OB_ (residues 109-1024) and p50_PDE_ (residues 255-270), absent in the crystal structure of the p50—p66_N_ complex^44^, face the p125 subunit but their weak density prevented *de novo* model building (Supplementary Figure 13).

It has been shown that DNA damage or replication stress induces the degradation of the p12 subunit, a 107-residue polypeptide of previously unknown structure, leading to a three-subunit enzyme with enhanced proofreading activity^19–21^. We identified p12 in the cryo-EM map as the cuboidal-shaped density stitched across the p125 and p50 subunits (Figure 1c), and we could model the p12 C-terminal portion spanning residues 42-107 (Figure 1d and 2e-f). The p12 N-terminus, which contains the PIP-box, is instead invisible, suggesting that it is flexible and projects away from the holoenzyme. The p12 fold is rather compact, with approximate dimensions of 30 Å × 20 Å × 20 Å, and consists of three short α-helices and loops interacting with the p125—p50 subcomplex, burying a total of ~1200 Å^2^ at the interface. The hydrophobic C-terminal loop of p12 (from Leu101 to Leu107) inserts like a hook into a pocket between the exonuclease domain and the CysB motif, establishing several contacts (Figure 2e-f). Additional polar interactions are observed between Gln51 on p12 helix α1 and Gly500 on the exonuclease domain, and between Thr62 on p12 helix α2 and Asp1068 in the ^CysB motif (Figure 2e). The interactions with the p50_OB_ are instead rather sparse (*e.g.*,^ Gln68 and Arg72 on p12 helix α2 interacting with Tyr388 on p50). Collectively, these results suggest a role for p12 as a scaffold stabilizing the catalytic module relative to the regulatory module of human Pol δ, enhancing the synthetic activity of the holoenzyme^18^ (Figure 2g). This stabilizing effect may regulate the small angular freedom between the two modules, which was observed in yeast Pol δ^23^ and suggested to facilitate the transfer of a mismatched primer from the catalytic to the exonuclease active site, a process critical for high-fidelity synthesis^45^ and predicted to involve a large rearrangement of the thumb and exonuclease domains^46,47^. Thus, degradation of p12 in response to DNA damage may slow down DNA synthesis, and the consequent increase of Pol δ flexibility may be required due to the higher rate of nucleotide misincorporation and frequent transfer of the primer between the pol and exonuclease sites.

### Interactions with DNA and implications for Pol δ activity

The catalytic module of human Pol δ is composed of N-terminal, palm, thumb and exonuclease domains (Figure 1d) and is structurally homologous to the catalytic module of yeast Pol3^48^. The positions of the P/T DNA and the incoming nucleotide in the polymerase active site are also analogous to those observed in the crystal structure of *Sc* Pol3–DNA–dCTP ternary complex^48^, with Watson-Crick base pairing between the terminal adenine in the template strand and the incoming dTTP (Figure 3, Inset 1). The density ascribed to the 5’-overhang of the template strand is compatible with three bound nucleotides that are sharply bent by a 90° angle relative to the duplex DNA and face the unbound protomer of PCNA (Figure 1d and 3). Therefore, the holoenzyme was captured in the act of synthesis. 22 base pairs (bp) of the DNA substrate could be modelled and thread through the PCNA hole in a central location, almost perpendicular to the ring plane (α_TILT_ ~ 4°) (Figure 3). The duplex DNA is in B-form and is kept in position by the thumb domain up to the 10th bp from the active site, while the remaining bases are stabilized by the constraint of the PCNA closed topology and interactions with basic residues in the clamp channel (Figure 3). In the absence of PCNA, the dsDNA portion below the thumb domain would be flexible and invisible in the map, as observed in the isolated *Sc* Pol δ holoenzyme bound to (P/T) DNA^23^. PCNA interacts with DNA via multiple polar residues approaching DNA phosphates within a coulombic interaction distance (< 6 Å) (Figure 3, Inset 2). Most of the interactions are established with one DNA strand, involving PCNA residues which follow the strand helix. A similar kind of electrostatic interactions were observed in the *lac* repressor binding to a non-specific DNA sequence^49^. The importance of the PCNA—DNA interactions for human Pol δ activity is supported by previous mutagenic studies, where a single mutation among PCNA basic residues K20, K77, K80, R149 and K217 at the sliding surface severely reduced Pol δ ability to incorporate an incoming nucleotide at the initiation of DNA synthesis^50^. Thus, such point mutations are expected to disrupt the observed pattern of interactions with DNA, resulting in a DNA tilt that is incompatible with DNA synthesis.

### Alternative Pol δ holoenzyme conformations reveal the importance of positioning the DNA in the central channel of PCNA

The local resolution map of the Pol δ holoenzyme showed reduced resolution of the PCNA component, particularly for the two unbound protomers, suggesting conformational heterogeneity in the complex (Supplementary Figures 1 and 4). In agreement, 3D classification identified different holoenzyme conformers in which the orientation of the PCNA ring plane varies in a range up to ~20° (Figure 4a and Supplementary Figure 5). Importantly, the tilting of PCNA disrupts the interactions involving residues 121-123 of the IDCL of PCNA and the CTD of p125 (Figure 2a) that are critical for the polymerization activity (Supplementary Figure 8), while the PIP-box interaction is maintained. The PCNA tilting also modulates the distance between a loop on the polymerase thumb domain and a loop on the PCNA front face (Figure 3b). Mutation of the highly conserved D41 residue on the PCNA loop (Supplementary Figure 8c) or K931A on the thumb loop (Supplementary Figure 10a) reduced the activity of Pol δ and severely compromised its processivity (Supplementary Figure 10b). This is consistent with previous findings showing the corresponding loop in *Sc* PCNA to be critical for enhancing the catalytic rate of the polymerase^15^. The contact between the two loops is visible in the lowest PCNA tilt conformer (Class 1, 4.3 Å resolution), which is the most populated (65%). However, the density at this contact is weak suggesting that the two loops interact in multiple orientations.

**Figure 5.**
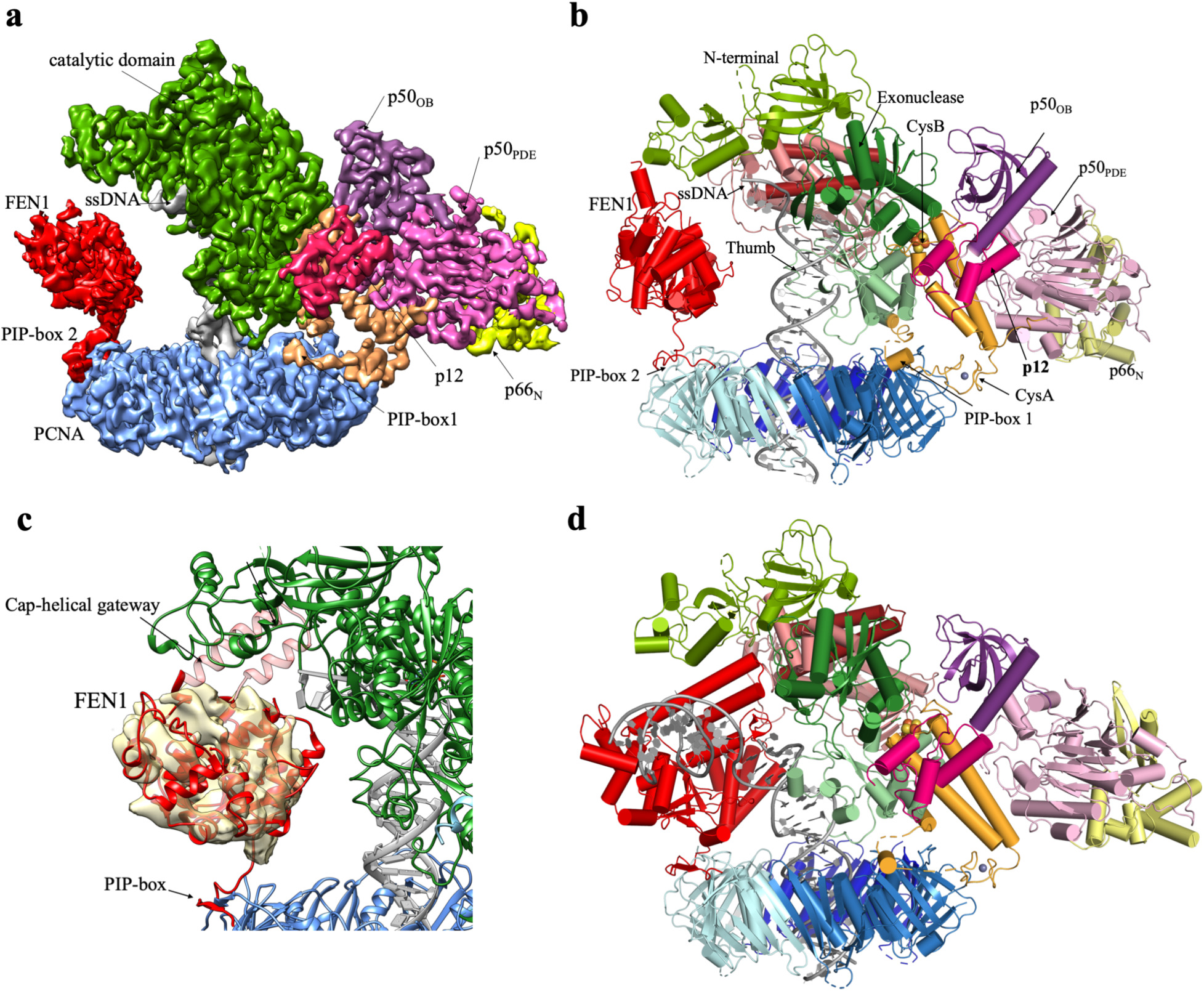
Cryo-EM structure of the Pol δ–DNA–PCNA–FEN1 complex. **a)** Cryo-EM map of the complex colored by domains. **b)** Complex model colored by domains. **c)** Close-up showing the map region around FEN1 filtered to 6 Å resolution. **d)** Proposed toolbelt model for Pol δ and FEN1 bound to PCNA processing an Okazaki fragment.

We propose that dynamic interactions among PCNA loops and both the thumb and the CTD of p125 may keep PCNA in a position that is competent for DNA polymerization. In the absence of such interactions (Class 2 and 3), the DNA tilts towards the PCNA inner rim and bends by ~10° to avoid clashing (Figure 3b and c). Interestingly, the DNA in Class 2 and 3 is translated vertically by one half of the major groove width relative to the processive holoenzyme (Figure 3d). As a consequence, the DNA strand mainly interacting with PCNA in the tilted conformers is the primer strand, while in the processive complex is the template strand. It is possible that this change in DNA positioning occurs frequently and disrupts the interactions of PCNA with the thumb and CTD. The binding via the PIP-box is integral for re-establishing the PCNA interactions with the thumb and CTD.

### Structure of the Pol δ—PCNA—DNA—FEN1 toolbelt and implications for Okazaki fragment maturation

The Pol δ holoenzyme architecture strongly suggests that other PCNA-interacting partners may bind to the PCNA monomers not occupied by the polymerase. Indeed, the strand displacement activity of Pol δ and 5’flap cleavage by FEN1 in yeast act processively through iterative 1-3 nt steps of nick translation to ultimately remove the RNA/DNA primer^16^. While these data indirectly support the “toolbelt” mechanism in nick translation, a polymerase—DNA—PCNA—FEN1 complex has not been observed before in eukaryotes. To test this hypothesis, we reconstituted a complex comprising human Pol δ, P/T DNA and dTTP, PCNA and FEN1, isolated the complex by gel filtration (Supplementary Figure 3), and analyzed it by cryo-EM (Supplementary Figures 14 and 15). We obtained a reconstruction at 4.0 Å resolution (Figure 5a) in which the mutual positions of Pol δ, DNA and PCNA are analogous to those of the processive Pol δ holoenzyme while FEN1 fills the space between the N-terminal and palm domains of the p125 subunit, where the ssDNA template enters the active site, and the unoccupied PIP-box site on PCNA. Previous crystallographic data showed that FEN1 interacts with PCNA in multiple orientations through a single C-terminal PIP-box connected to the FEN1 core by a flexible hinge encompassing residues 333-336^51^. Flexible-tethering of FEN1 to PCNA in the holoenzyme is suggested by the cryo-EM local resolution map showing lower resolution in the FEN1 region (6-9 Å, Supplementary Figure 14). Rigid-body fitting positions FEN1 in an upright configuration (Figure 5b-c) that is different from the three conformers observed in the FEN1—PCNA crystal structure, but that can be obtained by rotation and translation of any of these conformers via the flexible hinge. Density corresponding to the cap-helical gateway of FEN1, which forms a hole through which the 5’flap threads and is guided into the active site, is missing, suggesting that the cap-helical gateway is flexible and point outward from the polymerase. The reconstruction fitting a defined FEN1 conformer comprises ~20% of the particle dataset, while the reconstruction from the entire dataset shows no significant density at the FEN1 binding site (Supplementary Figure 4), suggesting that FEN1 may be absent from the complex in a major subset of particles or present in an ensemble of diverging orientations.

The Pol δ-DNA-PCNA—FEN1 complex structure reveals various aspects of the mechanism of maturation of Okazaki fragments. FEN1 sits across the template strand and in a proper orientation to bind the downstream duplex DNA when Pol δ encounters the previously synthesized Okazaki fragment (Figure 5b). Interestingly, the 90° bent angle of the template strand (Figure 3, Inset 1) is similar to that required for FEN1 activity^52^. This suggests that Pol δ may handoff an already bent nick junction to FEN1 and help in positioning the 5’ flap to thread into the cap-helical gateway (Figure 5d). However, handing off a bent DNA is not a must for FEN1 activity since FEN1 can actively bend the nick junction in diffusion-limited kinetics^53,54^. Nick translation reaction occurs at rates that are 10-fold faster than nick DNA product release by FEN1^16,54–56^. This suggests that Pol δ and FEN1 must be actively handing off their products during nick translation. Our structure suggests that the proximity of FEN1 to the template strand and the potential interaction between FEN1 and Pol δ may facilitate their products handoff during nick translation. The preferential binding of FEN1 to the more exposed PIP-box site on PCNA (Figure 5a) indicates that DNA Ligase 1 (Lig1) would be sterically excluded during nick translation reaction and that its binding requires Pol δ to move out of the way. The alternate conformers of PCNA may also facilitate the exposure of the third PIP-box site. This is consistent with the finding that Lig1 acts in a distributive manner during maturation of Okazaki fragments^16^. It is unclear how Pol δ—PCNA—FEN1 would signal the binding of Lig1. Interestingly, FEN1 releases the nick product in two steps, where it binds it briefly in a bent conformer that is followed by a lengthy binding in an extended conformer^54^. It is possible that the bent conformer is more compatible with Pol δ binding while the extended one sequesters the nick until Pol δ moves out of the way and Lig1 is recruited from solution. In yeast, acute depletion of Lig1 allows nick translation to proceed up to the dyad of the pre-assembled nucleosome, and the Okazaki fragment termini are enriched around known transcription factor binding sites^57^. Thus, nucleosomes and DNA binding factors may trigger the stalling of Pol δ and its dissociation from PCNA and/or DNA, allowing Lig1 to bind and seal the nick of a mature Okazaki fragment. Altogether, our findings pave the way to further structural and functional investigations of the complex roles of Pol δ in replicating the eukaryotic genome.

## Acknowledgements

This research was supported by King Abdullah University of Science and Technology through core funding (to S.M.H.), and by the Wellcome Trust (to A.D.B.). We acknowledge the Midlands Regional Cryo-EM facility at LISCB, major funder MRC.

## Author Contributions

M.T. purified Pol δ, M.T. and V.S.R. purified PCNA and its mutants, V.S.R. purified RFC, F.R. purified the initially used RFC using different protocol, and M.S.Z. and A.S. purified RPA; V.S.R. characterized the Fe-S; N.M. and F.J.B prepared PCNA for the cryo-EM analysis; C.L. prepared the cryo-EM samples; C.L., T.J.R., C.S. and A.D.B. analysed the cryo-EM data; C.L. and A.D.B built and refined the molecular models. M.T. performed the mutagenesis and functional assays. M.T., A.D.B. and S.M.H. designed the research and wrote the article. All authors analyzed the data, discussed the results and commented on the manuscript.

## Author Information

The maps of the Pol δ-DNA-PCNA complexes and Pol δ-DNA-PCNA—FEN1 complex have been deposited in the EMBD with accession codes EMD-10539, EMD-10080, EMD-10081, EMD-10082 and EMBD-10540 and the atomic models in the Protein Data Bank under accession codes PDB 6TNY, 6S1M, 6S1N, 6S1O and 6TNZ. The authors declare no competing financial interests. Correspondence and requests for materials should be addressed to S.M.H. or A.D.B.

## METHODS

### Plasmid Construction

MultiBac™ expression system (Geneva Biotech) was used to express human Pol δ in Sf9 insect cells. For this purpose, the genes encoding p125 (accession no. NP002682) and p50 (accession no. NP006221) were amplified and cloned into pACEBac1 Gent^+^ separately. The p50-encoding cassette along with promoter and terminator was excised with *I-Ceu1* and *BstX1* and ligated into p125 containing pACEBac1 linearized with *I-Ceu1*. After cloning wild-type p125 and p50, L1002A and F1005A (p125 PIP box) or K931A (p125) mutations were introduced by polymerase chain reaction (PCR) separately. The gene sequence encoding p66 (accession no. NP006582) was amplified and cloned into pIDC Cm^+^ while that of p12 (accession no. NP066996) was amplified along with an N-terminal 8xHis-tag and cloned into pIDS Spect^+^. After cloning wild-type p66 and p12, mutations of F462A and F463A (p66 PIP box) and I7A and Y11A (p12 PIP box) were carried out by PCR separately. Finally, the single transfer vectors with different subunit assemblies were generated using *Cre* recombinase according to the MultiBac™ expression system user manual and the resulting constructs were named hereafter as Pol δ, Pol δ-p125 PIP box mutant, Pol δ-p66 PIP box mutant, Pol δ-p12 PIP box mutant and Pol δ-K931A mutant. At the end, the recombinant transfer vector encoding all four Pol δ subunits was transformed into DH10MultiBac™ cells to transpose the gene expression cassettes into MultiBac baculoviral DNA and the resulting bacmid DNA was then isolated.

Full-length human PCNA (accession no. NM182649) was cloned in pETDuet-1 MCS1 (Novagen) Amp^+^ to obtain 6xHis N-terminally-tagged protein. After cloning wild-type PCNA (WT-PCNA), LA66A and 67E (hereafter named LA-PCNA) and D120A, L121A, D122A, and V123E (hereafter named DLDV-PCNA) and D41A (hereafter named D41A-PCNA) mutations were carried out by PCR. Human RPA clone (pET11d-tRPA) was a generous gift of Professor Marc S. Wold (University of Iowa College of Medicine, Iowa City, Iowa). A 6xHis-tag was introduced at the C-terminus of RPA1 subunit by PCR. The catalytically inactive FEN1-D181A was constructed as described previously^53^. Briefly, the gene sequence encoding WT human FEN1 was cloned via Gibson assembly into a pE-SUMO-pro expression vector (Lifesensors). An additional 6xHis-tag was introduced by PCR at the N-terminus of the fusion. D181A mutation was introduced by QuikChange II Site-Directed Mutagenesis Kit (Agilent).

For the human full-length RFC3 and RFC5 subunits, the expression vector pCDFK-RFC5/3 was a generous gift of Dr. Yuji Masuda^58^. pET *E. coli* expression vector with BioBrick polypromoter restriction sites (14-A) was a gift from Scott Gradia (Addgene plasmid # 48307)^59^. pCDFK-RFC5/3 was modified by inserting a 6xHis-tag at the N-terminus of RFC3. The sequences encoding RFC2, RFC4 and a truncated version of RFC1, with the first 550 N-terminal amino acids deleted, were codon optimized and synthesized by IDT as gBlocks. These three sequences were amplified by PCR with primers containing LICv2 sequences and cloned independently by Ligation Independent Cloning (LIC) into 14-A vectors. The three resulting expression cassettes were assembled together into one 14-A vector by BioBrick sub-cloning as instructed in the MacroBac manual^59^. This resulting co-expression plasmid is denoted from here onwards as p14A-ΔNRFC1/4/2.

### Protein expression and purification

For Pol δ, Pol δ-p125 PIP box mutant, Pol δ-p66 PIP box mutant, Pol δ-p12 PIP box mutant or Pol δ-K931A mutant expression, Sf9 cells were cultured in ESF 921 medium (Expression Systems). To prepare the baculovirus, bacmid DNA containing all four subunits was transfected into Sf9 cells using FuGENE® HD (Promega) according to manufacturer’s instructions. This baculovirus prep was then amplified twice to obtain a higher titer virus (P3 virus). The expression of Pol δ then proceeded by transfecting a 4L Sf9 suspension culture at a density of 2 × 10^6^ cells/ml with P3 virus. Cells were harvested at 72 hrs post-infection by centrifugation at 5,500 xg for 10 min and then re-suspended in 3 ml per 1 g of wet cells in lysis buffer [50 mM Tris-HCl (pH 7.5), 500 mM NaCl, 5mM β-Mercaptoethanol (BME), 0.2% NP-40, 1 mM PMSF, 5% (v/v) Glycerol and EDTA-free protease inhibitor cocktail tablet/50ml (Roche, UK)]. Cells were sonicated and debris was removed by centrifugation at 95,834 xg for 1 hr at 4 °C. The cleared lysate was then adjusted to 40 mM imidazole final concentration and directly loaded onto a HisTrap HP 5ml affinity column (GE Healthcare) pre-equilibrated with buffer A [50 mM Tris-HCl (pH 7.5), 500 mM NaCl, 5 mM BME and 5% Glycerol] containing 40 mM imidazole. After loading, the column was extensively washed first with 50 ml of buffer A containing 40 mM imidazole followed by washing with buffer A containing 100 mM imidazole to remove the nonspecifically-bound proteins. Finally, the column was washed with 50 ml of buffer B [50 mM Tris-HCl (pH 7.5), 50 mM NaCl, 40 mM Imidazole, 5 mM BME and 5% Glycerol] to reduce the salt concentration in preparation for next chromatographic step. The bound proteins were eluted with 50 ml linear gradient against buffer C [50 mM Tris-HCl (pH 7.5), 50 mM NaCl, 600 mM Imidazole, 5 mM BME and 5% Glycerol]. Fractions that contain all Pol δ subunits were combined and loaded onto an anion exchanger, Mono Q 5/50 GL (GE Healthcare) column pre-equilibrated with buffer D [50 mM Tris-HCl (pH 7.5), 50 mM NaCl, 5 mM BME and 5% Glycerol]. The column was then washed with 20 ml of buffer D containing 150 mM NaCl. Pol δ was eluted with a 20 ml of gradient from 150 mM NaCl to 1 M NaCl in 50 mM Tris-HCl (pH 7.5), 5 mM BME and 5% Glycerol. The fractions that contained all Pol δ subunits were combined and concentrated to 1 ml and loaded onto HiLoad 16/600 Superdex 200 pg (GE Healthcare) pre-equilibrated with gel filtration buffer [50 mM Tris-HCl (7.5), 200 mM NaCl, 1 mM DTT and 5% Glycerol]. Protein fractions were pooled, flash frozen and stored at −80 °C.

To express PCNA and its mutants, either WT-PCNA, LA-PCNA, DLDV-PCNA or D41A-PCNA plasmid was transformed into BL21 (DE3) *E. coli* cells and grown at 37 °C in 2YT media supplemented with ampicillin to an OD_600_ of 1.2. The cultures were then induced with 0.5 mM Isopropyl β-D-1-thiogalactopyranoside (IPTG) and continued to grow for 19 hrs at 16 °C. Cells were then harvested by centrifugation at 5,500 xg for 10 min. The resulting pellets were re-suspended in 3 ml per 1 g of wet cells in lysis buffer [50 mM Tris-HCl pH (7.5), 750 mM NaCl, 20 mM imidazole, 5 mM BME, 0.2% Nonidet P-40, 1 mM PMSF, 5% Glycerol and EDTA-free protease inhibitor cocktail tablet/50 ml]. The cells were lysed with 2 mg/ml of lysozyme at 4 °C for 30 min followed by sonication. Cell debris was removed by centrifugation (22,040 xg, 20 min, 4 °C) and the cleared supernatant was directly loaded onto HisTrap HP 5 ml affinity column pre-equilibrated with buffer A [50 mM Tris-HCl (pH 7.5), 500 mM NaCl, 20 mM Imidazole, 5 mM BME and 5% Glycerol]. The column was washed with 50 ml of buffer A followed by washing with 50 ml of buffer B [50 mM Tris-HCl (pH 7.5), 100 mM NaCl, 20 mM Imidazole, 5 mM BME and 5% Glycerol] to reduce the salt concentration. The bound PCNA, or its mutants, was eluted with 50 ml of gradient with buffer C [50 mM Tris-HCl (pH 7.5), 100 mM NaCl, 500 mM Imidazole, 5 mM BME and 5% Glycerol]. The eluents were pooled and directly loaded onto HiTrap Q HP 5 ml anion exchange column (GE Healthcare) pre-equilibrated with buffer D [50 mM Tris-HCl (pH 7.5), 100 mM NaCl, 5 mM BME and 5% Glycerol]. After loading, the column was washed with 50 ml of buffer D followed by elution with 50 ml gradient with buffer E [50 mM Tris-HCl (pH 7.5), 1 M NaCl, 5 mM BME and 5% Glycerol]. Eluents were pooled and concentrated to 1.5 ml and loaded onto HiLoad 16/600 Superdex 200 pg pre-equilibrated with gel filtration buffer [50 mM Tris-HCl (pH 7.5), 150 mM NaCl, 1 mM DTT and 5% Glycerol]. Protein fractions were pooled, flash frozen and stored at −80 °C. Recombinant PCNA used for the cryo-EM study was produced as described^60^.

Human RPA was expressed and purified as described previously^61^. Briefly, the plasmid was transformed into BL21 (DE3) *E. coli* cells and cultured in 2YT media at 37 °C. The expression was induced with 0.5 mM IPTG when the OD_600_ of the culture reached a value of 0.7. Buffer A is defined for all the rest of the RPA purification scheme as containing [50 mM Tris-HCl pH 7.5, 10% (v/v) glycerol and 10 mM β-mercaptoethanol (BME)]. The cells were collected and resuspended in lysis buffer [buffer A + 35 mM Imidazole and 750 mM NaCl]. RPA was first purified over a HisTrap HP 5 ml column pre-equilibrated with lysis buffer. The bound proteins were eluted with a linear gradient of buffer A + 250 mM Imidazole and 150 mM NaCl. From this step onward the purification followed the one described previously^62^ with few modifications as following. The collected fractions containing all the three RPA subunits were then loaded onto a HiTrap Blue 5 ml column (GE healthcare) pre-equilibrated with buffer A + 150 mM NaCl and washed extensively with buffer A + 1 M NaCl. The bound protein was then eluted with a linear gradient of buffer A + 1.5 M NaSCN. Finally, RPA was purified over a HiLoad Superdex-200 pg size exclusion column with buffer containing [50 mM Tris-HCl pH 7.5, 10% glycerol, 1 mM DTT, 0.1 EDTA]. The fractions containing pure stochiometric RPA were concentrated, flash frozen in liquid nitrogen and stored at −80 °C.

For the purification of RFC complex with an N-terminally truncated RFC1 (hereafter named as ΔNRFC), the two plasmids, p14A-ΔNRFC1/4/2 and pCDFK-RFC5/3, were co-transformed into BL21(DE3) *E. coli* cells and colonies were selected on agar plates containing two antibiotics (Kan + Amp). ΔNRFC was overproduced by growing the transformed cells in 10 L TB media containing the two antibiotics. Cells were grown at 20 °C to an OD_600_ of 0.8 and then expression was induced by the addition of 0.1 mM IPTG and continued at 15 °C for an additional 24 hrs. Cells were collected by centrifugation and re-suspended in buffer A [50 mM HEPES-NaOH (pH 7.5), 10 mM BME, 5% Glycerol and 1 mM PMSF] containing 500 mM NaCl and one EDTA-free protease inhibitor cocktail tablet/50 ml. Cells were lysed by lysozyme treatment and sonication and cleared by centrifugation at 95,834 xg for 45 min. The collected supernatant was adjusted to 10 mM imidazole and loaded onto a HisTrap HP 5 ml column. The salt concentration was lowered on-column to 250 mM NaCl and the protein complex was eluted with a linear gradient between Buffer A and Buffer A + 300 mM imidazole containing 250 mM NaCl. Fractions containing all the subunits of ΔNRFC were combined and loaded onto a HiTrap Heparin HP 1 ml column and eluted with linear gradient between Buffer A + 250 mM NaCl and Buffer A + 1 M NaCl. Fractions that contained all the ΔNRFC subunits were collected, concentrated and loaded onto HiLoad Superdex 16/600 200 pg gel filtration column equilibrated with storage buffer [50 mM HEPES-NaOH (pH 7.5), 300 mM NaCl, 1 mM DTT, 0.5 mM EDTA and 10% glycerol]. Fractions containing ΔNRFC with the correct stoichiometry were collected, concentrated, flash frozen and stored at −80 °C.

FEN1-D181A was expressed and purified as described previously^62^. Briefly, the plasmid was transformed into *E. coli* BL21 (DE3) cells and cultured in 2YT media. Expression was induced by the addition of 0.2 mM IPTG when the OD_600_ of the culture reached a value of 0.8. Cells were harvested and then lysed in buffer A [50 mM Tris–HCl pH 7.5, 5% (v/v) glycerol, 750 mM NaCl, 10 mM β-mercaptoethanol (BME) and 30 mM imidazole] by a combination of lysosome treatment and sonication. Purification was carried out by two sequential Ni-NTA columns, separated by SUMO protease cleavage. Proteins were then concentrated and further purified over HiLoad Superdex-75 pg size exclusion columns using a buffer containing [50 mM HEPES-KOH pH 7.5, 500 mM NaCl, 2 mM dithiothreitol (DTT), and 10% (v/v) glycerol]. Fractions containing pure D181A FEN1 were collected, flash frozen, and stored at −80 °C.

All protein concentrations were calculated by measuring their absorbance at 280 nm using the extinction coefficients calculated from the amino acid composition (Pol δ: 198140 M^−1^ cm^−1^, PCNA: 47790 M^−1^ cm^−1^, RPA: 87210 M^−1^ cm^−1^, FEN1: 22920 M^−1^ cm^−1^ and ΔNRFC: 157000 M^−1^ cm^−1^).

To check the trimeric structure of the PCNA mutants, a HiLoad 16/600 Superdex 200 pg gel filtration column was equilibrated with PCNA storage buffer. 2 mg of WT-PCNA, LA-PCNA and DLDV-PCNA were independently diluted in 250 μL of PCNA Storage Buffer. In parallel, a mixture containing 4 mg of Carbonic anhydrase (27 kDa) and Conalbumin (75 kDa) was prepared in 250 μL of PCNA storage buffer. Each sample was loaded onto the gel filtration column and ran for 1.5 column volumes at a flow-rate of 1 ml/min. The elution chromatograms were fit, using the cftool of MATLAB, to a single Gaussian in case of PCNA and to a sum of two Gaussians in case of the marker proteins. In both cases a linear baseline was added to the fit to correct for imperfect initialization of the UV_280_ reading. All three PCNA forms eluted between 71 and 72 ml (Supplementary Figure 7a-c). The two molecular markers eluted at ~77 ml and ~89 ml, respectively (Supplementary Figure 7d), at a later retention volume compared to PCNA. Since the PCNA monomer has a molecular weight of ~29 kDa, these results show that in all three forms, PCNA is larger than the largest of the two molecular markers of 75 kDa, therefore it must exist in its homotrimeric form.

### DNA substrates

M13mp18 single-stranded DNA (NEB N4040S) and Cy5 labelled HPLC purified primer (5’-/Cy5/TAA GGC CAG GGT TTT CCC AGT CAC G-3’) were annealed by mixing both templates in 1:1 molar ratio in T-50 buffer [50 mM Tris-HCl (pH 7.8), and 50 mM NaCl] and heating at 75 °C for 15 min that was followed by slow cooling to room temperature. For the substrate used in EM, a template strand (5’-CTGCACGAATTAAGCAATTCGTAATCATGGTCATAGCT-3’) was annealed to a primer containing a 3’ dideoxycytosine chain terminator (5’-AGCTATGACCATGATTACGAATTG[ddC]-3’) to form the P/TddC substrate. The strands were mixed in an equimolar ratio in the presence of 20 mM Tris pH 7.5 and 25 mM NaCl. They were then annealed by incubating at 92 °C for 2 minutes followed by slow cooling to room temperature overnight. All oligonucleotides were purchased from Sigma Aldrich.

### Primer extension assay

Primer extension assays were carried out in reaction buffer consisting of [40 mM Tris-HCl (pH 7.8), 5 mM MgCl_2_, 1 mM DTT, 0.2 mg/mL bovine serum albumin, 50 mM NaCl, 1 mM ATP and 500 μM each of dGTP, dATP, dTTP, and dCTP]. 5 nM of primed-M13 substrate was initially mixed with 150 nM PCNA (WT or mutants), 40 nM ΔNRFC, and 500 nM RPA on ice and then pre-incubated at 30 °C for 2 mins prior to the addition of different concentrations of Pol δ and further incubation for 10 mins except for D41A-PCNA where the reactions were incubated for 2.5 min; the reaction volume was 30 µl. Primer extension assays for Pol δ and its mutants in the presence of WT-PCNA were carried out at 30 °C for 2.5 min or for 5 min with or without 590 ng/ml heparin. Reactions with or without heparin were initiated with 500 μM dTTP. The reactions were terminated with 6 µl of stop buffer [180 mM NaOH, 6 mM EDTA and 7.5% Ficoll] followed by heating at 75 °C for 5 min and cooling on ice for 2 min. All products were resolved by running on 1% alkaline agarose gel for 20 hrs at 10 V and visualized using Typhoon Trio (GE Healthcare).

### UV-Vis analysis of Fe-S in purified human Pol δ

The absorption spectra of freshly purified Pol δ at four different concentrations were acquired at room temperature using a microplate spectrofluorometer (TECAN infinite M1000) in Pol δ storage buffer. Absorption spectra were collected from 260 to 600 nm in Corning 96-well UV-Plates. As per manufacturer’s instructions, 370 μL of each sample were used in order to generate an optical path length through the sample of 1 cm. Absorption spectra were also collected for background absorption using storage buffer as a blank. Criteria of 2-nm wavelength step size along with 100 flashes per step were used. The background absorption was then subtracted from the sample absorption spectra. For each of the four concentrations, four repeats of spectra were acquired. The final plotted spectra are the averages of the four individual repeats.

To quantify the presence of Fe-S cluster, the extinction coefficient at 410 nm was monitored^29,31,36,63,64^. It was previously indicated that the protein absorption peak, centered at ~280 nm, and the Fe-S absorption peak, centered at ~410 nm, are separated enough to be considered independent for quantification analysis^63,64^. Taking into account the Beer-Lambert law at each of these wavelengths, the following system of equations holds:

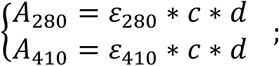

where A_280_ and A_410_ are the measured absorbances of the sample at 280 nm and 410 nm, ε_280_ and ε_410_ are the molar extinction coefficients of the sample at 280 nm and 410 nm, c is the concentration of the sample and d is the optical path length inside the sample. Given that for each sample the concentration and the optical path length are wavelength-independent, the above equations can be divided to remove the dependence on c and d, giving 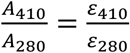. Multiplying both sides of this equation by ε_280_ we obtain the final equation of interest:

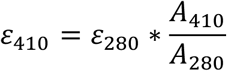

ε_280_ of Pol δ was calculated by Prot-Param^65^ to be 198140 M^−1^cm^−1^, based on its amino acids sequence and considering that all cysteines are in reduced form. Sixteen ε_410_ values were determined from the absorption spectra of the four repeats for each of the four Pol δ concentrations using the formula described above. The mean ε_410_ extinction coefficient together with its standard deviation (SD) were calculated from the sixteen values. It was previously established that each iron atom contributes with ~4000 M^−1^cm^−1^ to this extinction coefficient when coordinated inside an Fe-S cluster^63,64^. Therefore, the average and SD of the number of iron atoms per Pol δ are obtained by dividing the mean and SD of ε_410_ by 4000 M^−1^cm^−1^.

### Cryo-EM grid preparation and data collection

For all complexes, UltrAuFoil® R1.2/1.3 Au 300 grids were glow discharged for 5 minutes at 40 mA on a Quorum Gloqube glow-discharge unit, then covered with a layer of graphene oxide (Sigma) prior to application of sample. For preparation of the Pol δ complex without FEN1, the P/TddC substrate was mixed, in order, with Pol δ, PCNA and dTTP. Final concentrations were 0.84 μM Pol δ and PCNA trimer, 2.1 μM DNA and 20 μM dTTP. The buffer this was performed in comprised 25 mM HEPES-KOH (pH 7.5), 100 mM potassium acetate, 10 mM calcium chloride, 0.02% NP-40, 0.4 mM biotin and 1 mM DTT. 3 μl of this sample were applied to the grid, blotted for 3 seconds at blot force 10 and plunge frozen into liquid ethane using a Vitrobot Mark IV (FEI Thermo Fisher), which was set at 4 °C and 100% humidity. For the Pol δ–DNA–PCNA–FEN1 complex, a 40 μl inject containing 3.75 μM P/TddC, 1.5 μM Pol δ, 1.5 μM PCNA trimer, 20 μM dTTP, and 3 μM FEN1, was loaded onto a Superdex 200 increase 3.2/300 column (GE Life Sciences) equilibrated with the same buffer as above (Supplementary Fig. 3). 3 μl of a fraction corresponding to the first peak was applied to a grid, prepared in the same way as above for data collection. Cryo-EM data for all samples were collected on a Thermo Fisher Scientific Titan Krios G3 transmission electron microscope at the LISCB at Leicester University. Electron micrographs for the complex with multiple PCNA conformers were recorded using a Falcon III direct electron detector (FEI Thermo Fisher) at a dose rate of 0.7 e-/pix/sec for 60 seconds and a calibrated pixel size of 1.08 Å. These data were collected using a Volta phase plate with EPU 1.9, and focusing performed at every hole using a nominal value of −0.6 μm. For the toolbelt structure, electron micrographs were recorded using a K3 direct electron detector (Gatan Inc.) at a dose rate of 11 e-/pix/sec and a calibrated pixel size of 0.87 Å. Focusing was performed over a range between −2.3 and −1.1 μm, in 0.7 μm intervals.

### Cryo-EM image processing

Pre-processing of the processive Pol δ complex (Dataset 1) was performed as follows: movie stacks were corrected for beam-induced motion and then integrated using MotionCor2^66^. All frames were retained and a patch alignment of 4 × 4 was used. Contrast transfer function (CTF) parameters for each micrograph were estimated by Gctf^67^. Integrated movies were inspected with Relion-3.0^68^ for further image processing (3575 movies). Particle picking was performed in an automated mode using the Laplacian-of-Gaussian (LoG) filter implemented in Relion-3.0. All further image processing was performed in Relion-3.0. Particle extraction was carried out from micrographs using a box size of 360 pixels (pixel size: 1.08 Å/pixel). An initial dataset of 2.8×10^6^ particles was cleaned by 2D classification followed by 3D classification with alignment. 3D refinement and several rounds of polishing and per-particle CTF refinement yielded a 4.09 Å structure of the Pol δ–PCNA complex. Masked refinement of the Pol δ and PCNA components yielded reconstructions at 3.88 and 6.70 Å, respectively. Classification of the Pol δ holoenzyme showing PCNA in different orientations was performed as follows: particles previously aligned on the Pol δ component using masked refinement were subsequently 3D-classified without alignment. Five 3D classes were generated with populations corresponding to 3, 4, 6, 22 and 65%. These classes were aligned on the Pol δ component and sorted for the tilting of the PCNA ring plane relative to the polymerase. 3D refinement was performed on the 3D classes corresponding to populations 65, 22 and 6%, and yielded reconstructions at 4.3, 4.9 and 8.1 Å resolution. Pre-processing of the Pol δ–DNA– PCNA–FEN1 complex data (Dataset 2) was performed as follows: movie stacks were imported in super resolution mode, then corrected for beam-induced motion and integrated using Relion’s own implementation, using a binning factor of 2. All frames were retained and a patch alignment of 5 × 5 was used. Contrast transfer function (CTF) parameters for each micrograph were estimated by CTFFIND-4.1^69^. Integrated movies were inspected with Relion-3.1 for further image processing (5071 movies). Particle picking was performed in an automated mode using the Laplacian-of-Gaussian (LoG) filter implemented in Relion-3.1.

Further image processing was performed in Relion-3.1, taking two separate paths to generate the reconstructions with and without visible FEN1. The former was processed as follows: Particle extraction was carried out from micrographs using a box size of 400 pixels (pixel size: 0.87 Å/pixel). An initial dataset of 2.5×10^6^ particles was cleaned by 2D classification followed by 3D classification with alignment. 3D refinement and several rounds of polishing and per-particle CTF refinement yielded a 4.05 Å structure of the Pol δ–PCNA-FEN1 toolbelt complex, with clear density corresponding to FEN1. The data without visible FEN1 was processed as follows: An initial dataset of 2.5×10^6^ particles was cleaned by 2D classification followed by 3D classification with alignment. 3D refinement yielded a map of the holoenzyme at 4.6 Å resolution. Further 3D classification was then performed without alignment, followed by 3D refinement and several rounds of polishing and CTF refinement, yielding a 3.0 Å structure of the holoenzyme.

### Molecular modelling

The homology model for the catalytic domain of human Pol δ was built with Phyre2^70^ based on the crystal structure of the catalytic domain of *Saccharomyces cerevisiae* (*Sc*) Pol δ (PDB entry 3IAY)^48^, rigid-body docked and jiggle-fitted using Coot^71^. For enhanced fitting precision, the chain of the human Pol δ catalytic domain was partitioned into 12 segments (residues 2-83; 84-161; 162-288; 289-352; 352-371; 372-408; 409-496; 497-575; 576-638; 639-756; 757-873; 874-908) corresponding to the homologous segments in *Sc* Pol δ undergoing TLS vibrational motion as determined from the *Sc* Pol δ crystal structure^48^ with TLS Motion Determination (TLSMD)^62^, and the segments were fitted in the cryo-EM map as individual rigid bodies. The homology model for the zinc finger in the CTD of Pol δ was built with HHpred^72^, based on the crystal structure of Spt4/5NGN heterodimer complex from *Methanococcus jannaschii* (PDB entry 3LPE)^73^, and edited with Coot. Main-chain tracing was aided by ARP/wARP^74^. The homology model for CysB motif of human Pol δ was built with Phyre2^70^ based on the cryo-EM structure of *Saccharomyces cerevisiae* (*Sc*) Pol δ (PDB entry 6P1H)^23^. Modelling of the DNA and dTTP in the catalytic site of Pol δ was guided by the crystal structure of the *Sc* Pol δ—DNA—dCTP complex^48^. DNA was built with Coot and 3D-DART^75^. The model of the p12 subunit of Pol δ was built using ARP/wARP^74^, and edited with Coot. Rigid-body docking of the human p50—p66N complex (PDB entry 3E0J)^44^ and PCNA (PDB entry 1U76)^27^ was performed in UCSF Chimera^76^. The model of the processive Pol δ complex was subjected to real-space refinement in Phenix^77^. FEN1 core (chain Y in PDB entry 1UL1^51^) was rigid body fitted in the EM map of the Pol δ–DNA–PCNA–FEN1 complex using Chimera and FEN1 hinge (residues 333-336) was edited with Coot to have the C-terminal PIP-box anchored to its binding site on PCNA. The Pol δ component in Conformers 1, 2 and 3 was built from that of the processive complex, while PCNA was rigid-body fitted in the corresponding maps, and the region of the CTD connecting the PIP-box to the CysA motif was edited with Coot.

