## Supplementary Data for "Structure of the processive human Pol δ holoenzyme"

[1]Leicester Institute of Structural & Chemical Biology and Department of Molecular & Cell Biology, University of Leicester, Lancaster Rd, Leicester LE1 7HB, UK. [2]Division of Biological and Environmental Sciences and Engineering, King Abdullah University of Science and Technology, Thuwal 23955, Saudi Arabia. [3]CIC bioGUNE, Parque Tecnológico de Bizkaia Edificio 800, 48160 Derio, Spain. [4]IKERBASQUE, Basque Foundation for Science, Bilbao, Spain.

* These authors contributed equally to the work

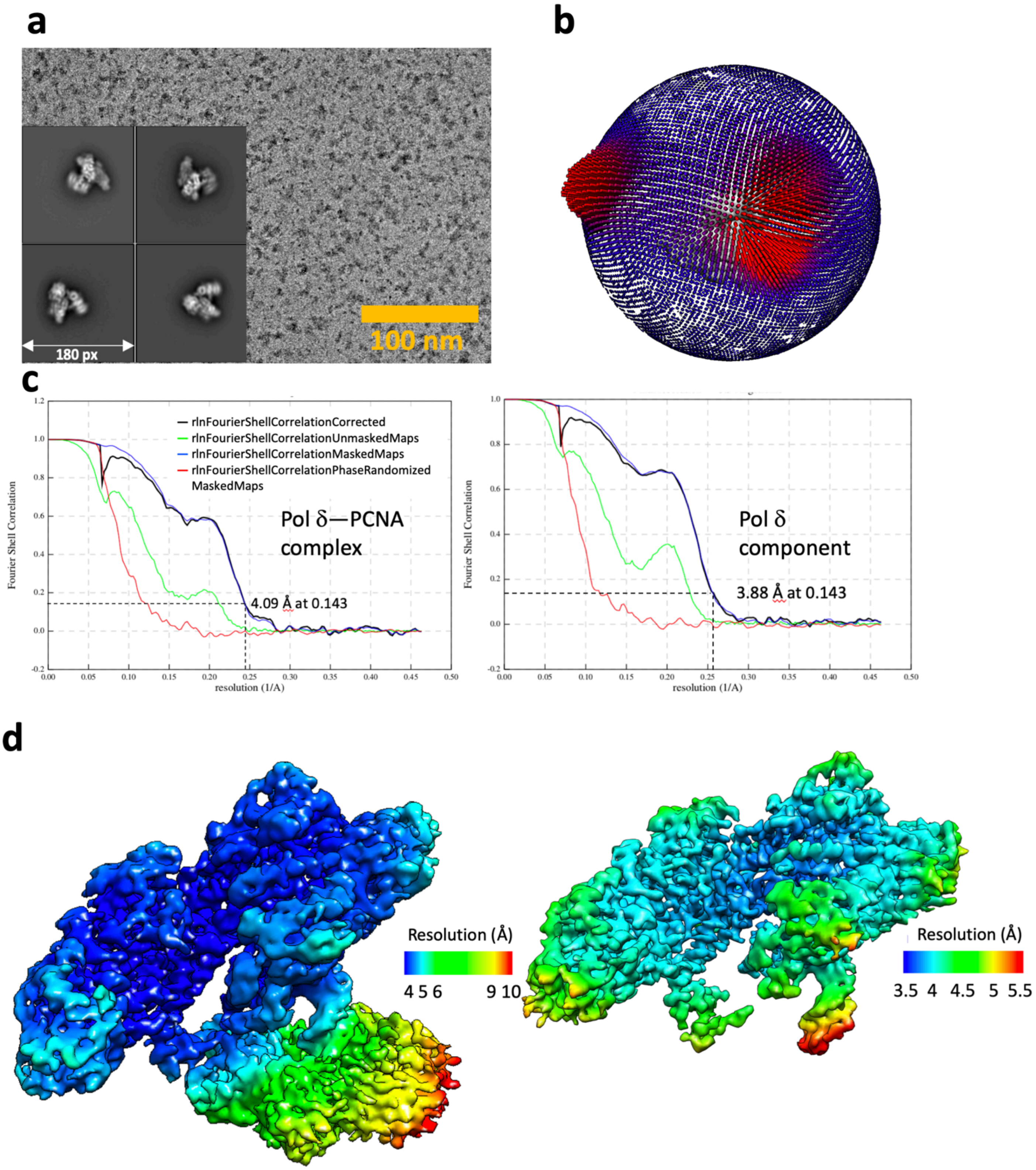

**Supplementary Figure 1.** Cryo-EM of the processive Pol δ holoenzyme (Dataset 1). **a)** Electron micrograph (aligned sum) acquired using a Volta phase plate on a Falcon 3EC direct electron detector in counting mode, and representative 2D class averages. **b)** Angular distribution of projections. **c)** Gold-standard Fourier shell correlation for the reconstruction of the full complex or the polymerase component after focussed refinement, and resolution estimation using the 0.143 criterion. **d)** Cryo-EM maps of the full complex (left) and polymerase component after focussed refinement (right) color-coded by local resolution.

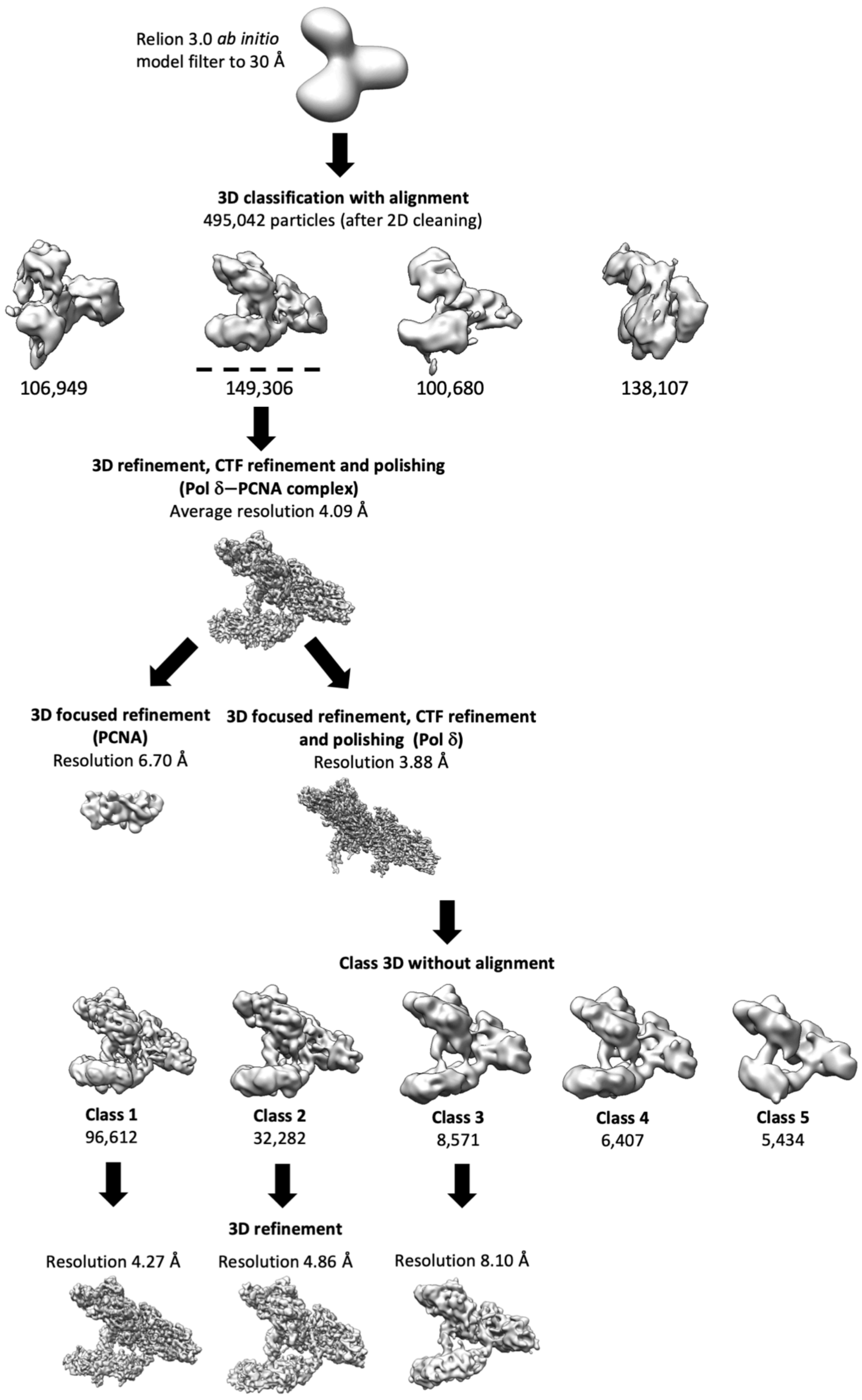

**Supplementary Figure 2.** Overview of image processing of Pol δ holoenzyme with different PCNA conformers (Dataset 1).

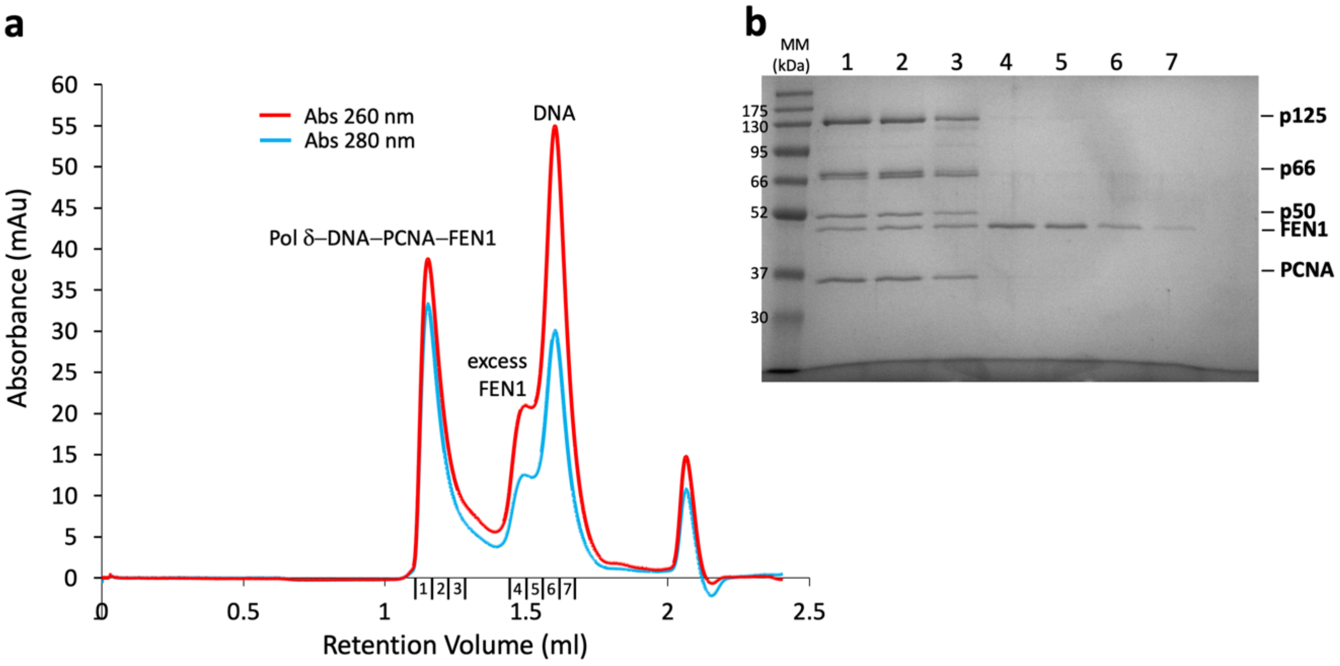

**Supplementary Figure 3.** Sample separation of the Pol δ-DNA-PCNA-FEN1 complex. **a)** Gel filtration chromatography of the reconstituted Pol δ-DNA-PCNA-FEN1 complex. **b)** The numbered peak fractions in a) were analysed by SDS-PAGE (lanes 1–7). Proteins corresponding to the bands are labeled on the right. Molecular weight standards are shown on the left.

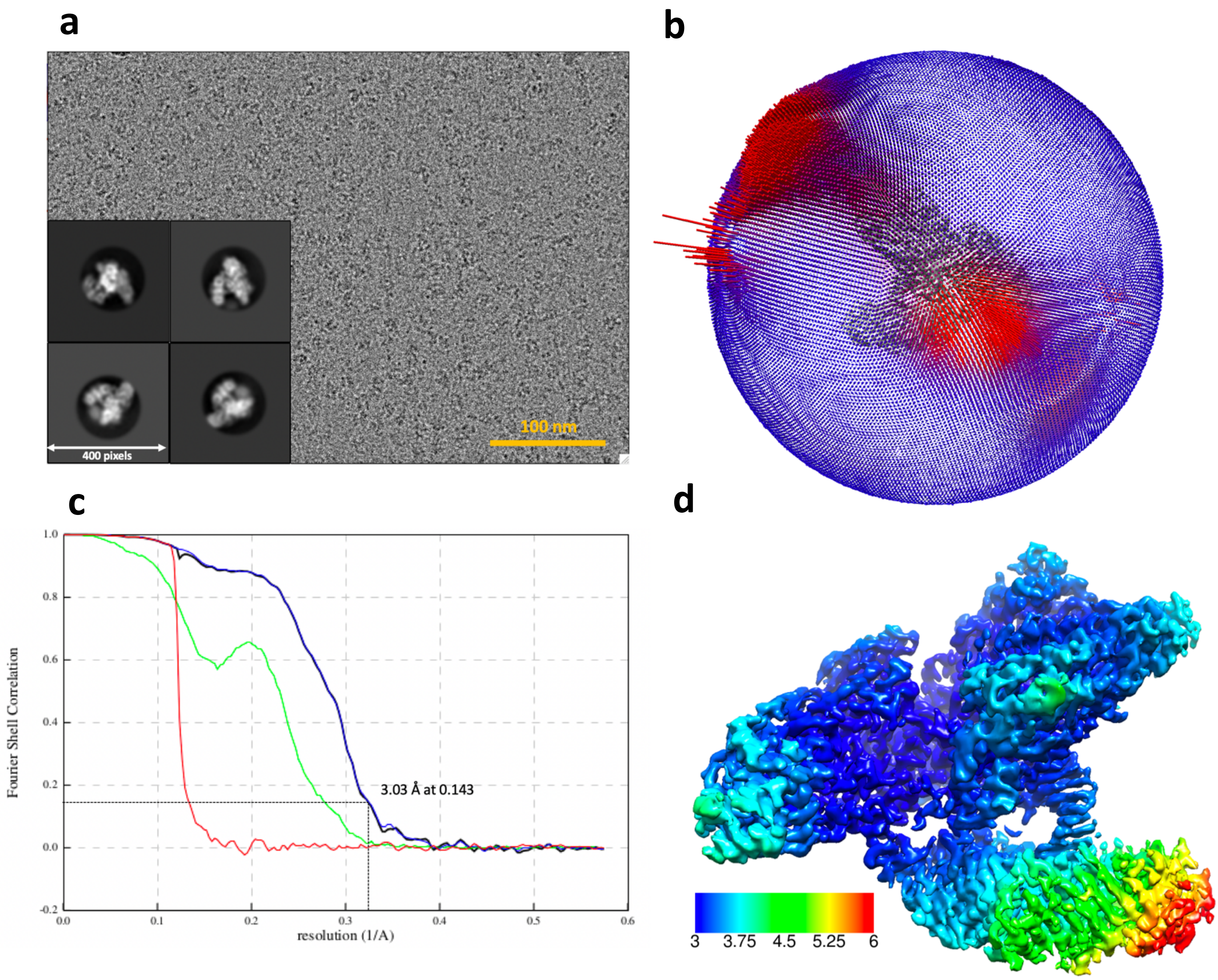

**Supplementary Figure 4.** Cryo-EM of the processive Pol δ holoenzyme (Dataset 2). **a)** Electron micrograph (aligned sum) acquired on a K3 direct electron detector in super resolution mode, and representative 2D class averages. **b)** Angular distribution of projections. **c)** Gold-standard Fourier shell correlation for the complex, and resolution estimation using the 0.143 criterion. **d)** Cryo-EM map of the complex, color-coded by local resolution.

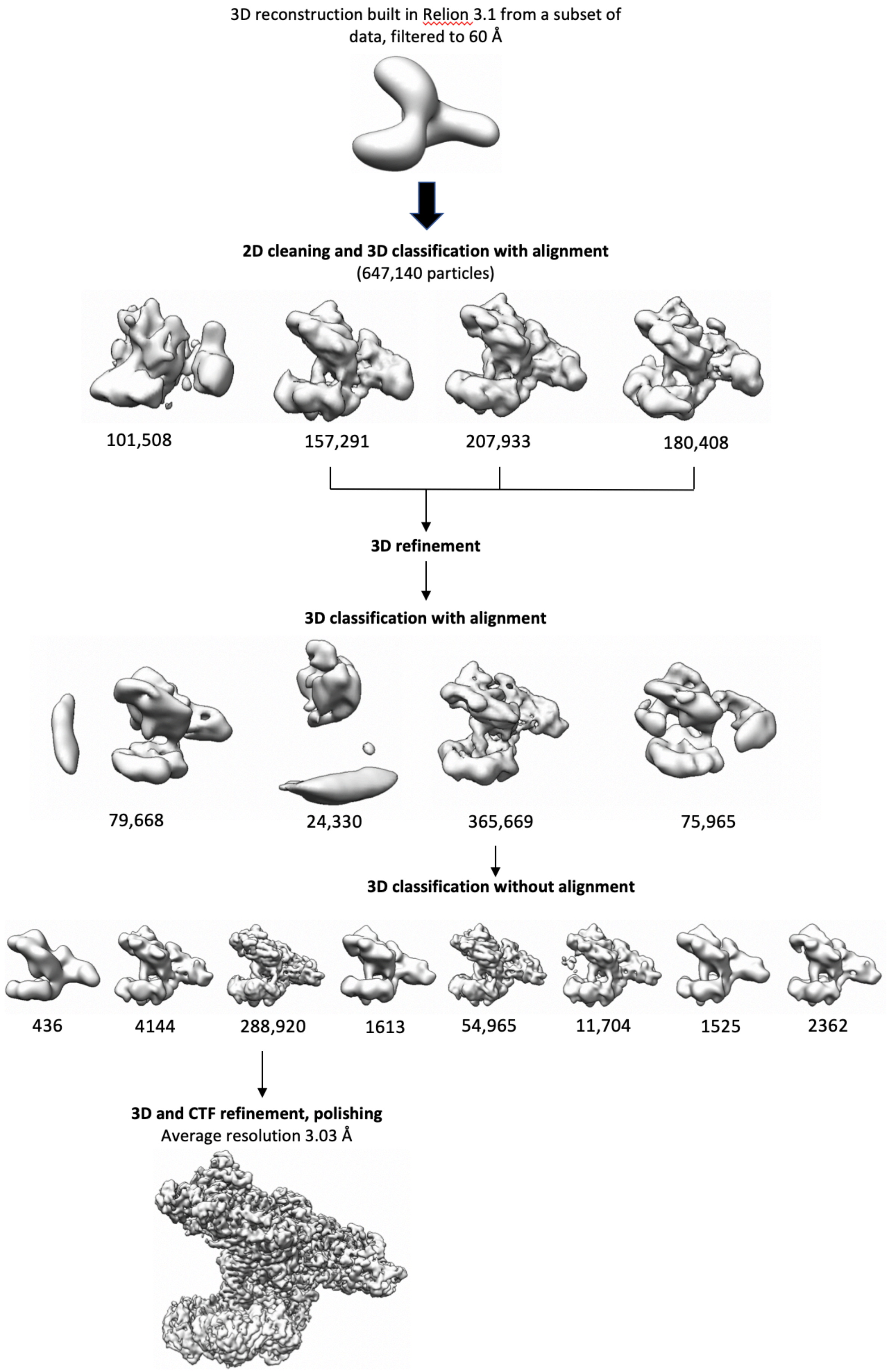

**Supplementary Figure 5.** Overview of image processing of the processive Pol δ holoenzyme (Dataset 2).

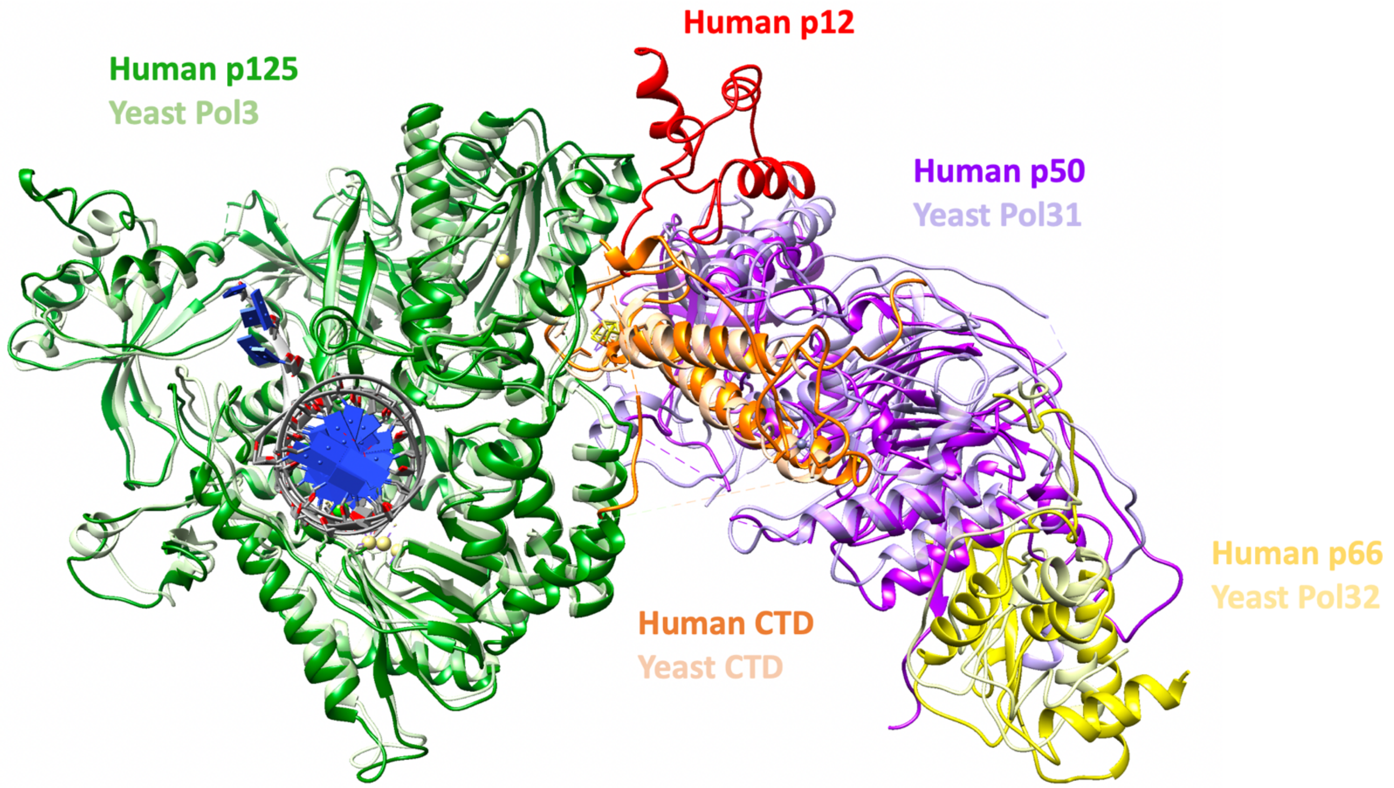

**Supplementary Figure 6.** Processive human Pol δ holoenzyme overlaid with the *S. cerevisiae* homologue (PDB: 6P1H). For clarity, PCNA was removed from the processive Pol δ holoenzyme structure.

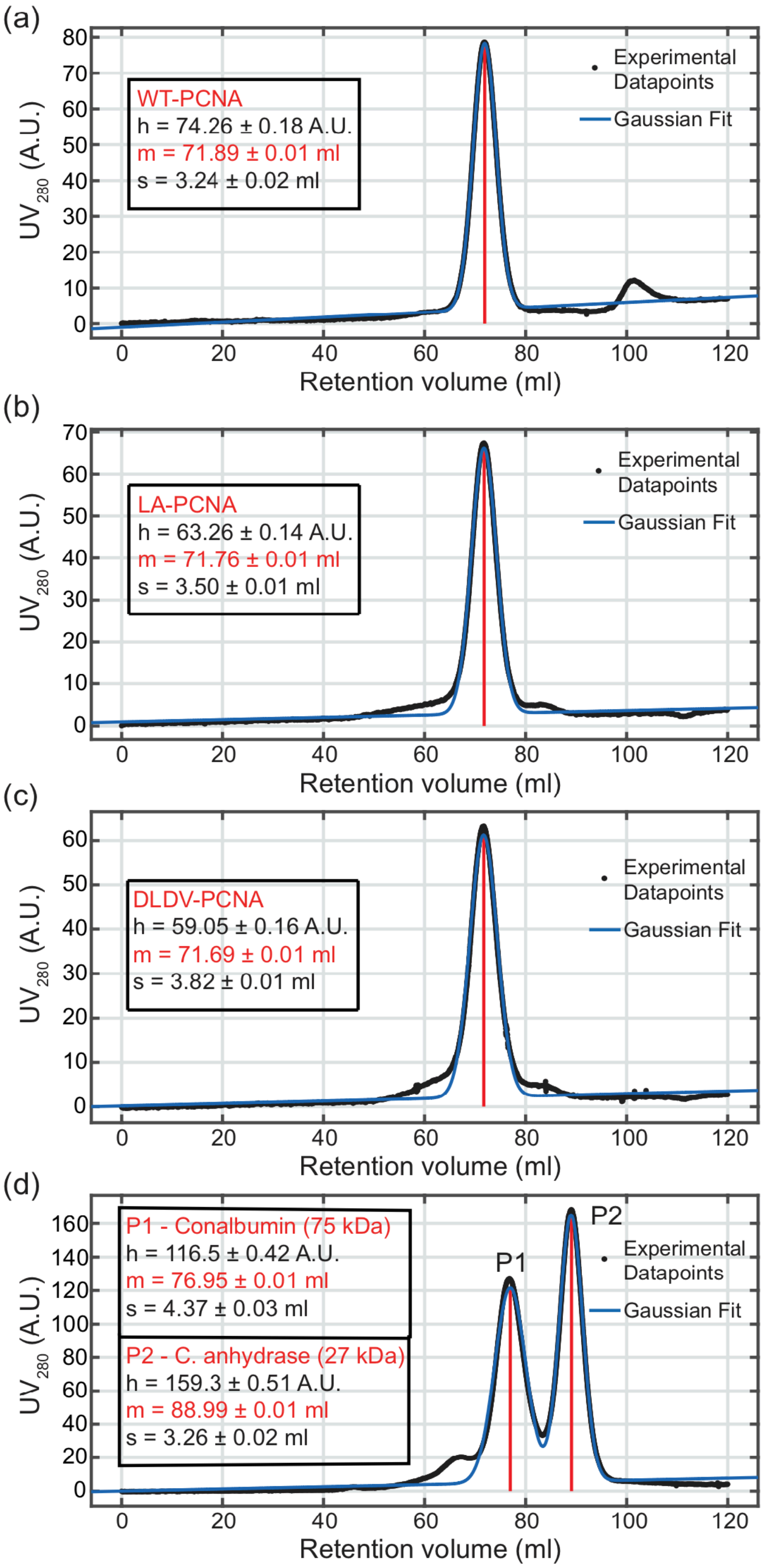

**Supplementary Figure 7.** Gel filtration profiles of WT PCNA and mutants PCNA. The fitted elution profiles are shown for the following proteins: **a)** WT-PCNA, **b)** LA-PCNA, **c)** DLDV-PCNA, **d)** Molecular weight markers Carbonic anhydrase (27 kDa) and Conalbumin (75 kDa). Fittings of the elution peaks of the purified WT PCNA and mutant PCNAs and comparison to the elution of molecular markers. The values of the Gaussian fitting parameters are recorded in the inset tables together with their 95% confidence interval. The vertical red lines represent the positions of the maxima of the elution peaks.

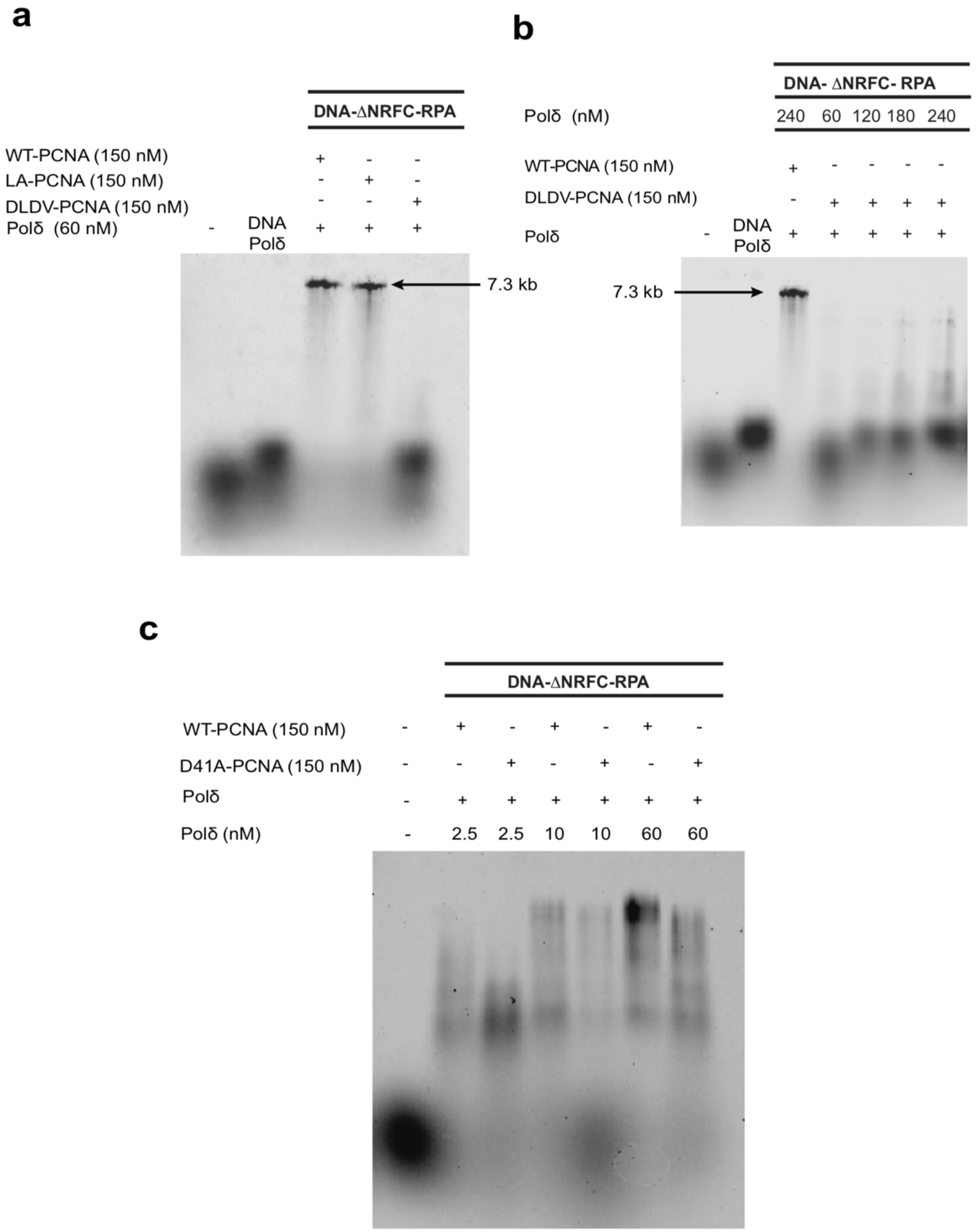

**Supplementary Figure 8.** Primer extension assays with Pol δ and PCNA mutants. **a)** M13mp18 DNA primer extension assay with WT-PCNA, LA-PCNA and DLDV-PCNA mutants as indicated. Reactions were carried out at 30 °C for 10 min **b)** M13mp18 DNA primer extension assay with different concentrations of Pol **δ** using DLDV-PCNA mutant. Reactions were carried out at 30 °C for 10 min. **c)** M13mp18 DNA primer extension assay with different concentrations of Pol δ using D41A-PCNA mutant. Reactions were carried out at 30 °C for 2.30 min. Products were separated on 1% alkaline agarose gel at 10 V for 20 hrs.

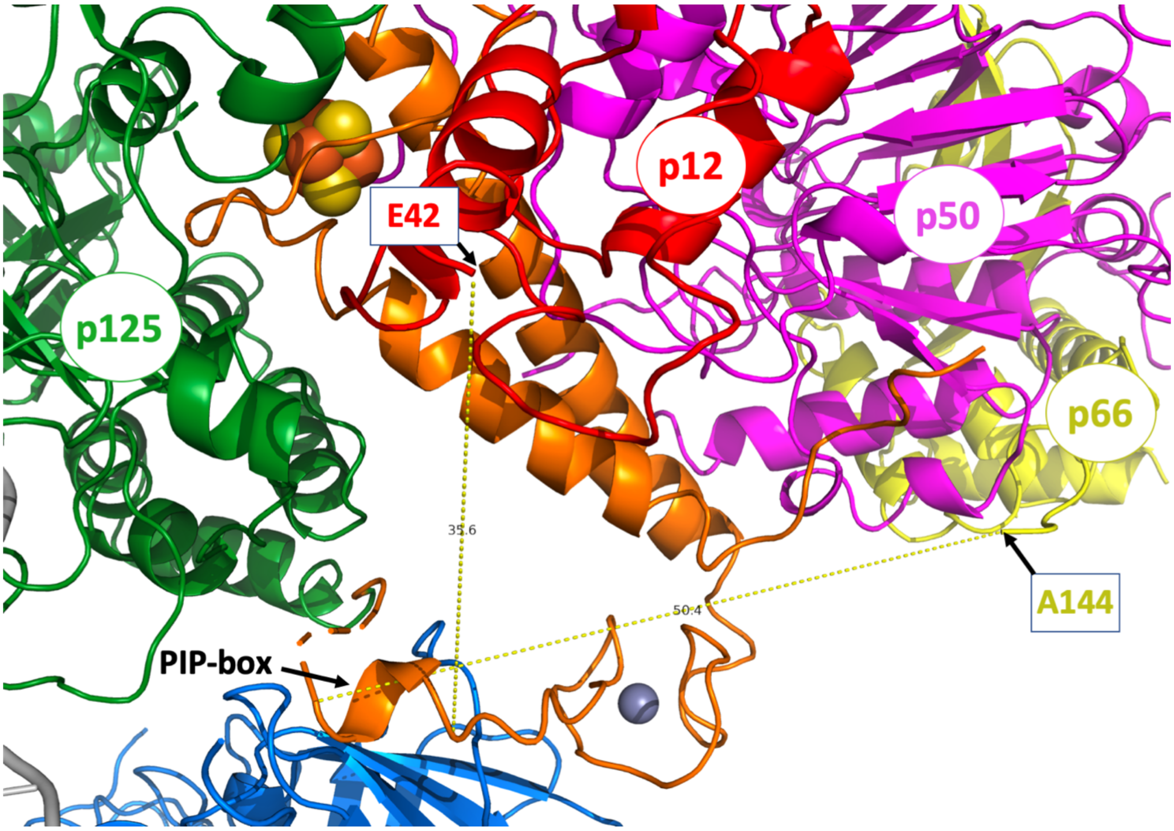

**Supplementary Figure 9.** Distance between the last visible residue of p66 (A144, 308 residues from the p66 PIP-box) and the N-terminus of the CTD PIP-box is ~50 Å. The distance between the first visible residue of p12 (E42, 31 residues from the p12 PIP-box) and the C-terminus of the PIP-box is ~36 Å.

**a**

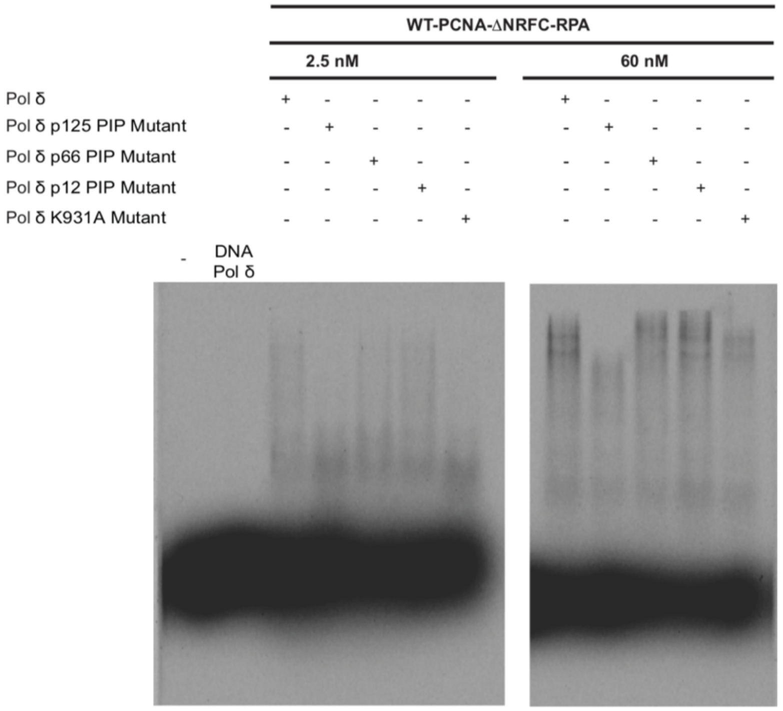

**b**

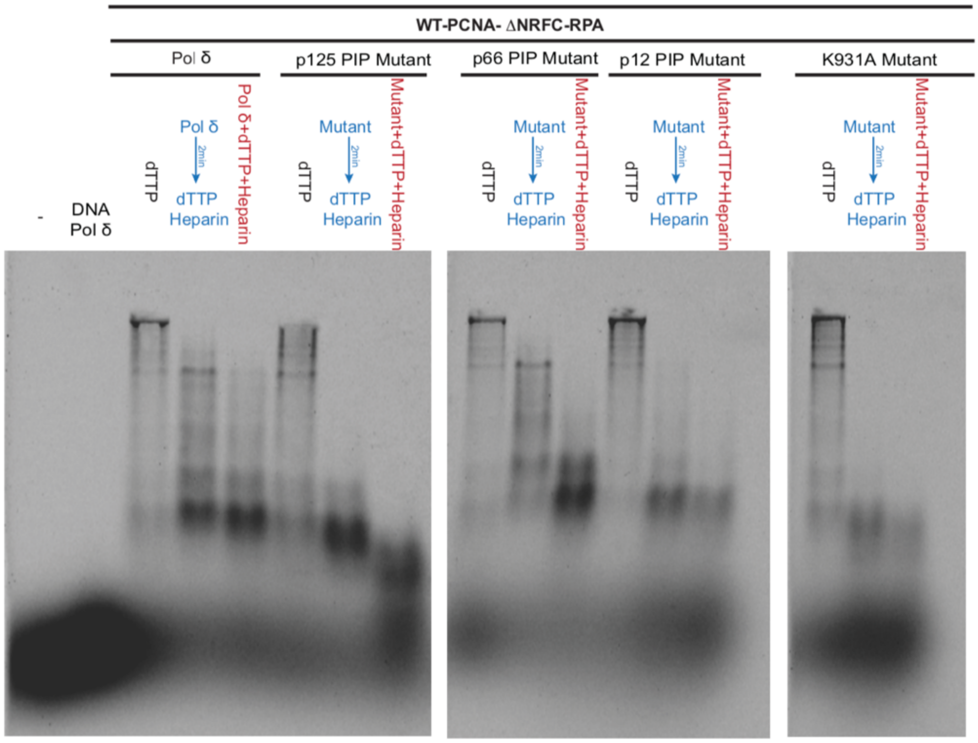

**Supplementary Figure 10.** Primer extension and processivity assays with Pol δ and Pol δ mutants. **a)** M13mp18 DNA primer extension assay with different concentrations of Pol δ, Pol δ p125 PIP mutant, Pol δ p66 PIP mutant, Pol δ p12 PIP mutant and Pol δ K931A mutant as indicated. Reactions were carried out at 30 °C for 2.30 min. **b)** Processivity of Pol δ, Pol δ p125 PIP mutant, Pol δ p66 PIP mutant, Pol δ p12 PIP mutant and Pol δ K931A mutant with or without 500 ng/ml heparin. Reactions were initiated with 500 μM dTTP with or without heparin. All reactions were carried out at 30 °C for 5 min and products were separated on 1% alkaline agarose gel at 10 V for 20 hrs.

**a**

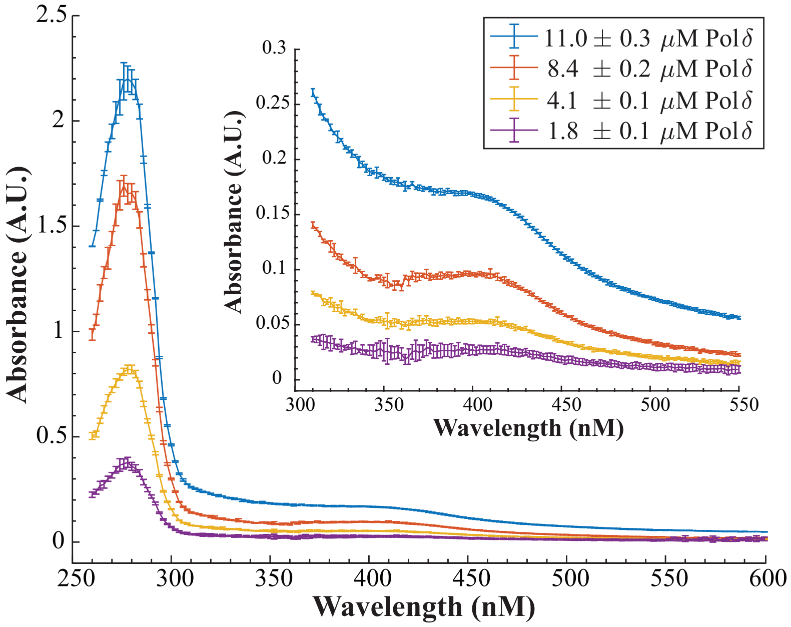

**b**

| | | Pol δ Concentration (μM) | | | |
|---|---|---|---|---|---|
| | **Repeat** | **11.0 ± 0.3** | **8.4 ± 0.2** | **4.1 ± 0.1** | **1.8 ± 0.1** |
| **$A_{280}$ (A.U.)** | 1 | 2.2347 | 1.6241 | 0.7995 | 0.3447 |
| | 2 | 2.1299 | 1.6975 | 0.8297 | 0.3754 |
| | 3 | 2.2351 | 1.6169 | 0.8057 | 0.3421 |
| | 4 | 2.137 | 1.6994 | 0.8429 | 0.3763 |
| **$A_{410}$ (A.U.)** | 1 | 0.1633 | 0.0963 | 0.0558 | 0.0327 |
| | 2 | 0.1624 | 0.0937 | 0.0526 | 0.0249 |
| | 3 | 0.1661 | 0.1024 | 0.0513 | 0.0257 |
| | 4 | 0.1652 | 0.0934 | 0.0522 | 0.0253 |
| **$\varepsilon_{410}=\varepsilon_{410}{}^{*}A_{410}/A_{280}$ ($M^{-1}cm^{-1}$)** | 1 | 14479.02 | 11748.59 | 13828.91 | 18796.57 |
| | 2 | 15107.72 | 10937.09 | 12561.36 | 13142.48 |
| | 3 | 14724.64 | 12548.42 | 12615.84 | 14885.12 |
| | 4 | 15317.14 | 10922.02 | 12270.62 | 13321.66 |

**Supplementary Figure 11.** Characterization of the FeS cluster in Pol δ. **a**) UV-Vis absorption spectra in the region 260-600 nm of Pol δ at four concentrations. The spectra show a broad absorption band which has a peak at ~410 nm that scales with the Pol δ concentration. The inset shows a zoomed in view of the spectra that focuses on the 410 nm broad absorption band. The spectra were acquired as described in the Methods section. Each data point in each spectrum represents the average of four measurements. The error bars represent the standard deviation (SD) of the four measurements. **b)** Absorbance measurements and calculations of Pol δ extinction coefficient at 410 nm and the average number of iron atoms per protein. Values are calculated as described in Methods section from four measurements per each Pol δ concentration. Mean $\varepsilon_{410}$= 13575.45 ± 1980.83 $M^{-1}.cm^{-1}$ and [Fe]/[ Pol δ] = 3.4 ± 0.5.

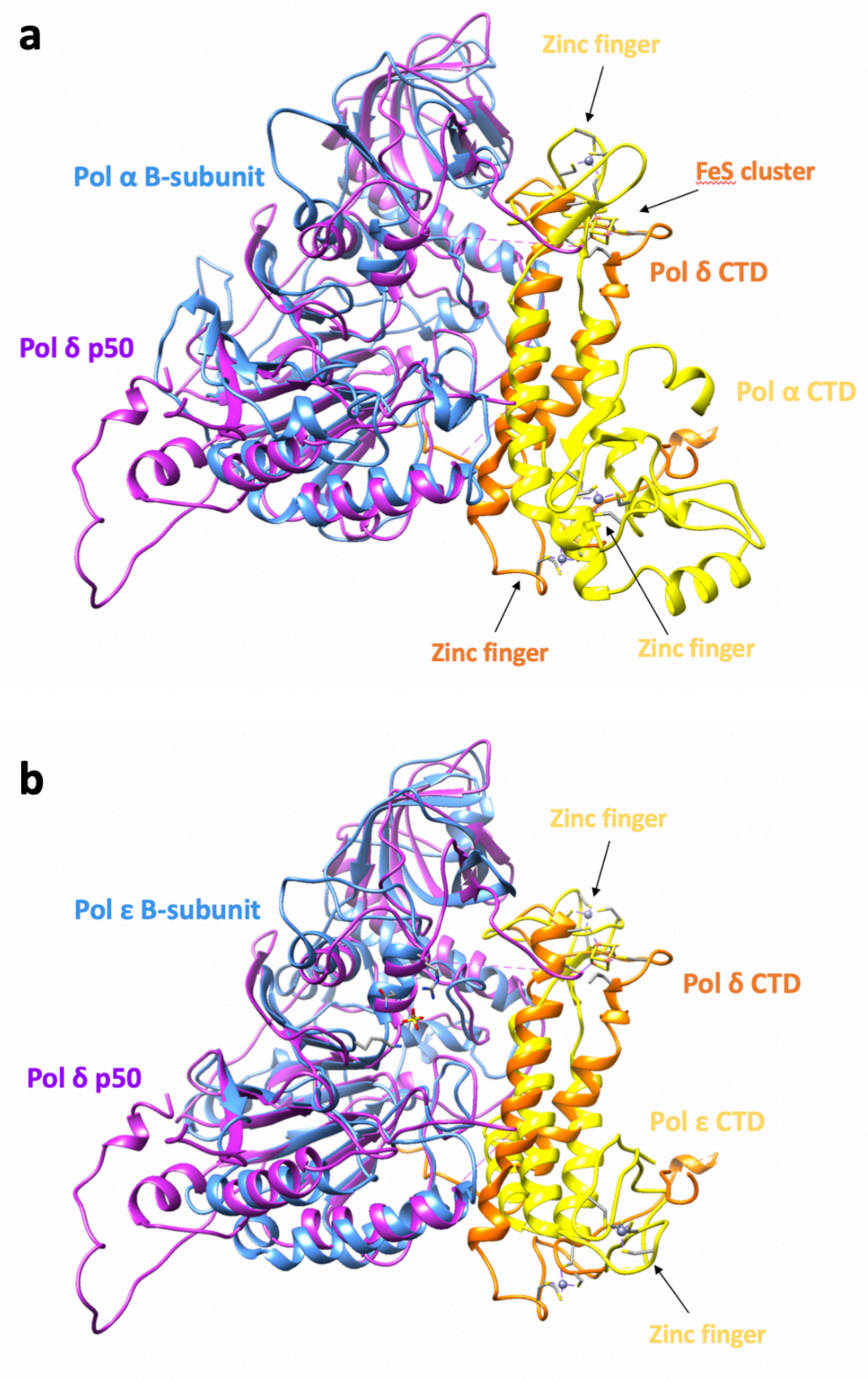

**Supplementary Figure 12.** Orientation of the CTD relative to the B-subunit in Pol δ, α and ε. a) Comparison of the B-subunit–CTD complex of Pol δ with Pol α. b) Comparison of the B-subunit–CTD complex of Pol δ with Pol ε. The cryo-EM model of Pol δ was aligned to that of Pol α (PDB entry 4Y97) or Pol ε (PDB entry 5VBN) using the B-subunit. The metal binding sites in the CTDs are labelled. Models are shown in ribbon representation. Zinc ion and Fe-S centre are shown as a sphere and cube, respectively.

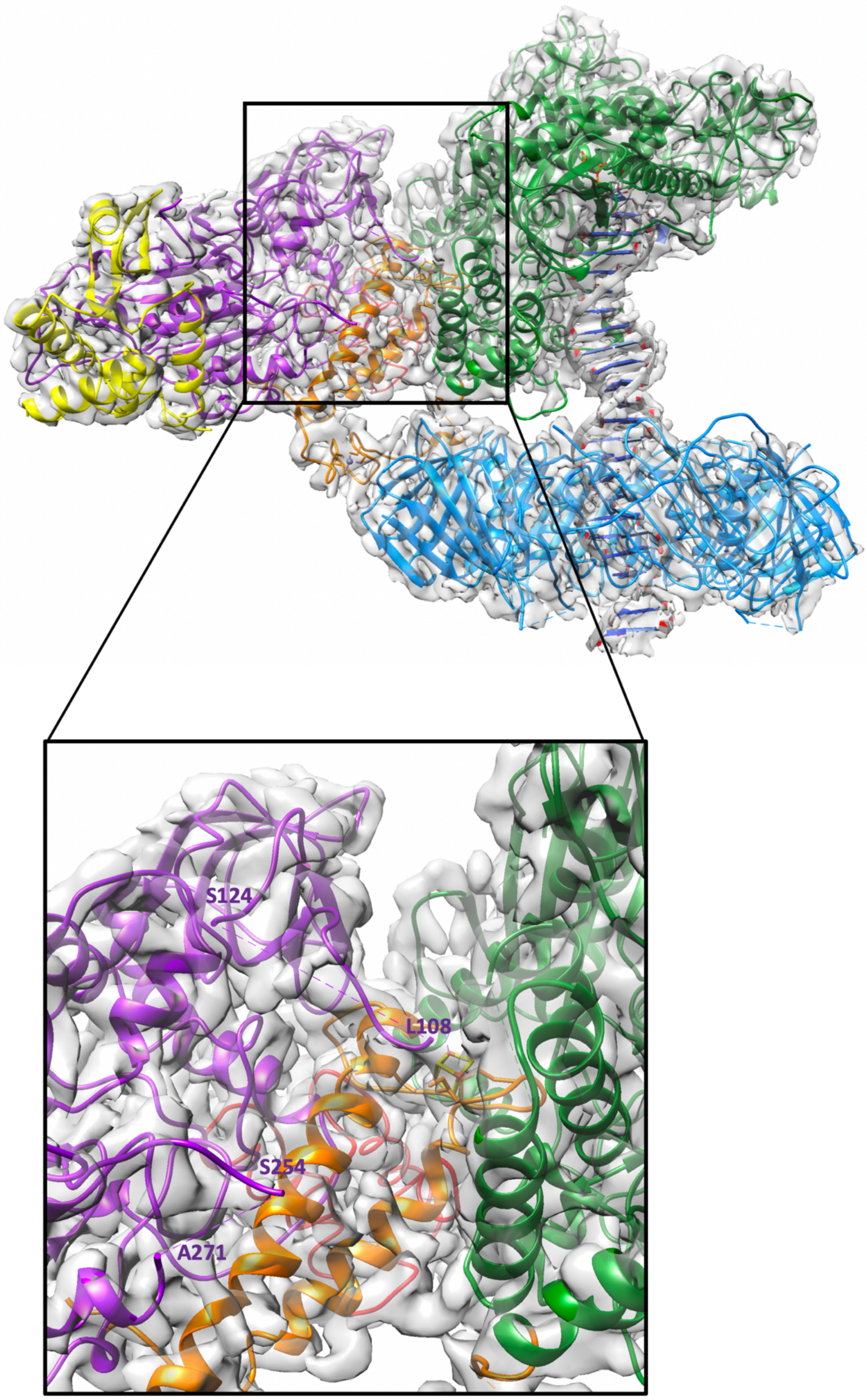

**Supplementary Figure 13.** Inset (bottom) from the model of the processive holoenzyme (top) showing the unmodeled loops of p50 (255-270 and 109-123) which face the catalytic subunit.

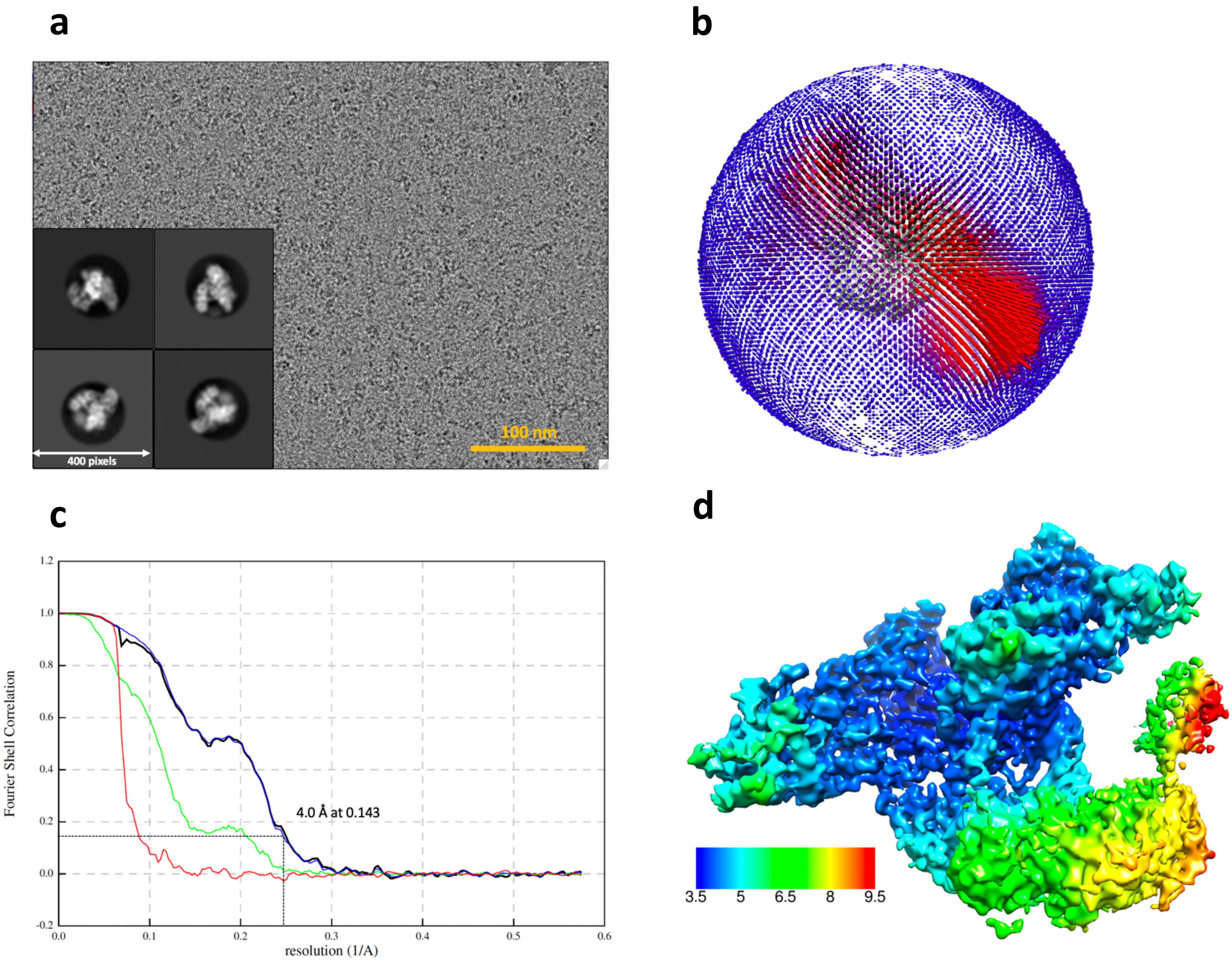

**Supplementary Figure 14.** Cryo-EM of the Pol δ-DNA-PCNA-FEN1 toolbelt complex. **a)** Electron micrograph (aligned sum) acquired on a K3 direct electron detector in super resolution mode, and representative 2D class averages. **b)** Angular distribution of projections. **c)** Gold-standard Fourier shell correlation for the complex, and resolution estimation using the 0.143 criterion. **d)** Cryo-EM map of the complex, color-coded by local resolution.

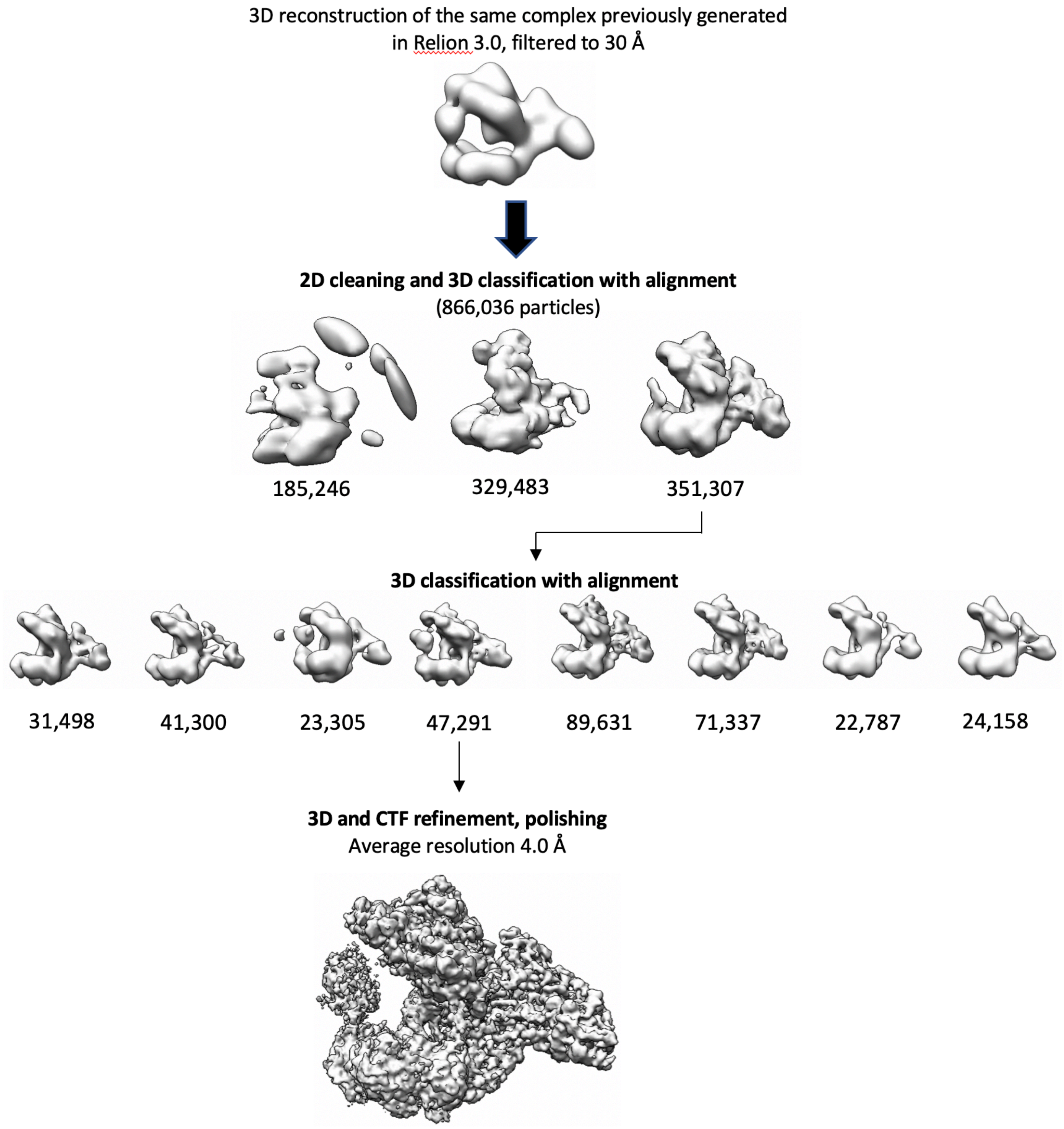

**Supplementary Figure 15.** Overview of image processing of the Pol δ-DNA-PCNA-FEN1 complex.

**Supplementary Table 1 |** Cryo-EM data collection, refinement and validation statistics

| | Processive Polδ-DNA-PCNA complex (EMD- 10539) (PDB 6TNY) | Polδ-DNA-PCNA-FEN1 complex (EMDB-10540) (PDB 6TNZ) | Polδ-DNA-PCNA complex Conformer 1 (EMDB- 10080) (PDB 6S1M) | Polδ-DNA-PCNA complex Conformer 2 (EMDB- 10081) (PDB 6S1N) | Polδ-DNA-PCNA complex Conformer 3 (EMDB- 10082) (PDB 6S1O) |
|---|---|---|---|---|---|
| **Data collection and processing** | | | | | |
| Magnification | 105,000 | 105,000 | 75,000 | 75,000 | 75,000 |
| Voltage (kV) | 300 | 300 | 300 | 300 | 300 |
| Electron exposure (e–/Å$^2$) | 44 | 44 | 35 | 35 | 35 |
| Defocus range (μm) | -1.1 to -2.3 | -1.1 to -2.3 | -0.4 to -0.6 | -0.4 to -0.6 | -0.4 to -0.6 |
| Pixel size (Å) | 0.87 | 0.87 | 1.08 | 1.08 | 1.08 |
| Symmetry imposed | C1 | C1 | C1 | C1 | C1 |
| Initial particle images (no.) | 2,512,757 | 2,512,757 | 2,821,297 | 2,821,297 | 2,821,297 |
| Final particle images (no.) | 288,920 | 47,291 | 96,612 | 32,282 | 8571 |
| Map resolution (Å) | 3.03 | 4.05 | 4.27 | 4.86 | 8.10 |
| FSC threshold | 0.143 | 0.143 | 0.143 | 0.143 | 0.143 |
| Map resolution range (Å) | 2.84-15.5 | 3.68-10.0 | 4.02-10.7 | 4.42-15.4 | 9.19-24.4 |
| **Refinement** | | | | | |
| Initial model used (PDB code) | | | | | |
| Model resolution (Å) | 3.03 | 4.05 | 4.27 | 4.86 | 8.10 |
| FSC threshold | 0.143 | 0.143 | 0.143 | 0.143 | 0.143 |
| Model resolution range (Å) | 2.84-15.5 | 3.68-10.0 | 4.02-10.7 | 4.42-15.4 | 9.19-24.4 |
| Map sharpening *B* factor (Å$^2$) | -5.32 | 6.05 | 5.64 | 7.45 | 6.18 |
| Model composition | | | | | |
| Non-hydrogen atoms | 19,728 | 22,010 | 19728 | 19728 | 19728 |
| Protein residues | 2399 | 2705 | 2399 | 2399 | 2399 |
| Nucleotide residues | 50 | 50 | 50 | 50 | 50 |
| Ligands | 3 | 3 | 3 | 3 | 3 |
| *B* factors (Å$^2$) | | | | | |
| Protein | 63.41 | 65.03 | 50.28 | | |
| Nucleotide | 149.74 | 149.74 | 163.98 | | |
| Ligand | 31.02 | 31.02 | 31.02 | | |
| R.m.s. deviations | | | | | |
| Bond lengths (Å) | 0.007 | 0.007 | 0.008 | 0.009 | 0.009 |
| Bond angles (°) | 1.093 | 1.130 | 1.215 | 1.322 | 1.312 |
| Validation | | | | | |
| MolProbity score | 1.87 | 2.31 | 2.24 | 2.35 | 2.30 |
| Clashscore | 7.94 | 13.59 | 13.13 | 14.59 | 14.59 |
| Poor rotamers (%) | 0.83 | 1.57 | 1.46 | 1.96 | 1.70 |
| Ramachandran plot | | | | | |
| Favored (%) | 93.24 | 91.08 | 91.93 | 92.75 | 92.79 |
| Allowed (%) | 6.42 | 7.84 | 7.22 | 6.72 | 6.68 |
| Disallowed (%) | 0.34 | 1.09 | 0.84 | 0.53 | 0.53 |
